# Phosphate resupply differentially impacts the shoot and root proteomes of *Arabidopsis thaliana* seedlings

**DOI:** 10.1101/2025.06.28.661392

**Authors:** Milena A. Smith, Lauren E. Grubb, Kirsten H. Benidickson, Devang Mehta, William C. Plaxton, R. Glen Uhrig

**Affiliations:** Department of Biology, Queen’s University, Kingston, Ontario, Canada; Department of Biological Sciences, University of Alberta, Alberta, Canada; Department of Biosystems, KU Leuven, Leuven, Belgium; Leuven Plant Institute, KU Leuven, Leuven, Belgium; Department of Biomedical and Molecular Sciences, Queen’s University, Kingston, Ontario, Canada; Department of Biochemistry, University of Alberta, Edmonton, Alberta, Canada

## Abstract

Phosphate (Pi) is an essential macronutrient for plant development that is often limited in soil environments. Plants have evolved myriad dynamic biochemical, physiological, and morphological adaptations to cope with nutritional Pi deficiency, collectively known as the Pi starvation response (PSR). While many components of the PSR have been well-characterized, much less is known about how metabolic homeostasis is re-established upon Pi resupply, particularly with respect to tissue- and time-specific adaptations. Here, we applied label-free quantitative proteomics to quantify protein-level changes in *Arabidopsis thaliana* shoots and roots following Pi resupply after prolonged Pi deprivation, quantifying a total of ∼2,700 differentially abundant proteins (DAPs). Sampling at early (1 h) and late (48 h) time-points, we captured a combination of immediate signaling and metabolic responses, along with longer-term recovery processes. Early responses prioritized metabolic adjustments to restore Pi pools via enhanced glycolysis and energy production, followed by later shifts toward anabolism, including nucleotide production, membrane remodeling, and protein synthesis. Several key enzymes, including ALTERNATIVE OXIDASE 1A, FRUCTOSE-BISPHOSPHATE ALDOLASE 5, and subunits of PHOTOSYSTEM I exhibited unique tissue-specific and time-dependent regulation. Overall, our findings reveal dynamic temporal phases of metabolic reprogramming during recovery from Pi starvation, and identify candidate proteins involved in orchestrating this transition, ultimately identifying potential targets for enhancing Pi uptake- and use-efficiency in crops. While hydroponic liquid culture enabled precise control of Pi availability, soil responses may be further influenced by heterogeneity and other root interactions.

## 1 INTRODUCTION

Phosphorus (P) is an essential plant macronutrient required for fundamental biochemical, metabolic, and physiological processes, with root epidermal cells directly assimilating this mineral from the soil as orthophosphate (H_2_PO_4_^-^; Pi) via plasmalemma Pi transporters (PHTs) (Dissanayaka *et al*., 2021). The role of Pi in plants is multifaceted. It serves as a central structural component of important biomolecules, including nucleic acids, sugar phosphates, and phospholipids, and is also required for key processes such as photosynthesis (i.e. photophosphorylation, and triose-P exchange across chloroplast envelope) and respiration (i.e. ATP synthase of inner mitochondrial membrane). Pi also serves as an allosteric activator or inhibitor of several key regulatory enzymes of central plant metabolism, while controlling the function of many proteins via its covalent attachment to specific amino acid residues of phosphoproteins. Despite Pi’s significance, most soil-Pi levels are far below those required to sustain optimal crop growth. Pi concentrations in the soil solution rarely exceed 10 μM owing to Pi’s propensity to precipitate as insoluble metal cation-Pi complexes (Lambers and Plaxton, 2015). Furthermore, a significant proportion of soil-Pi is trapped in organic phosphorous (Po) compounds (i.e. decaying biomatter), which must be mineralized prior to plant uptake (Dissanayaka *et al*., 2021). Suboptimal levels of soluble Pi in soils have resulted in agricultural applications of massive quantities of unsustainable and polluting Pi fertilizers that are sourced from rapidly depleting ‘rock-Pi’ reserves (Dissanayaka *et al.,* 2021; Hinsinger, 2001; Blackwell *et al*., 2019). Specifically, Pi runoff from fertilized fields contaminates aquatic and marine ecosystems, triggering harmful algal and cyanobacterial blooms and eutrophication (Lambers and Plaxton, 2015). Given the inefficiency of Pi fertilizers and scarcity of rock- Pi reserves, precise agricultural solutions are urgently needed to reduce our over-reliance on exogenous Pi fertilizer application. By studying the adaptations of Pi-deficient (-Pi) plants, we may uncover potential biological targets for engineering Pi-efficient crop varieties.

Given their sessile nature, -Pi plants have evolved a variety of physiological, morphological, and biochemical adaptations that function to enhance Pi-acquisition and -use efficiency, collectively known as the Pi starvation response (PSR). The PSR is regulated by a combination of Pi-starvation inducible (PSI) genes, along with various post-transcriptional and post-translational controls, ultimately increasing Pi-acquisition and -use efficiency (Dissanayaka *et al*., 2021). Given the low mobility of Pi, -Pi plants expand their root networks (i.e., increased lateral root formation and root hair density) to enhance foraging and thereby increase Pi-uptake (Dissanayaka *et al.,* 2021). Furthermore, PSI high-affinity PHTs belonging to the PHT1 family actively transport Pi across the plasma membrane of root cells against a steep concentration gradient (Dissanayaka *et al*., 2021). To mobilize soil Pi, roots of -Pi plants also excrete organic acid anions (particularly malate and citrate) and Pi-scavenging enzymes (e.g., nucleases and acid phosphatases (APases)) to solubilize Pi bound to metal cations and trapped in Po substrates, respectively (Dissanayaka *et al*., 2021). Plants acclimate to limiting Pi though a range of Pi-sensing and scavenging mechanisms; notably, members of the PHT2-5 families regulate Pi exchange and mobilization between subcellular compartments, whereas secreted and vacuolar purple APases (PAPs) respectively hydrolyze Pi from extracellular (i.e. cell wall, apoplast, and rhizosphere) and intracellular Pi-monoesters and anhydrides (Dissanayaka *et al*., 2021). Additionally, -Pi plants exhibit remarkable metabolic flexibility by upregulating alternative enzymes to ‘bypass’ various adenylate- and Pi-dependent reactions of central plant metabolism. For example, inorganic pyrophosphate (PPi)-dependent enzymes, such as PPi-dependent phosphofructokinase, UDP-glucose pyrophosphorylase, and the tonoplast H^+^-PPiase have been reported to be upregulated under -Pi conditions, shifting flux away from ATP-dependent pathways when cytosolic ATP, but not PPi pools become significantly depleted (Palma *et al*., 2000; Dissanayaka *et al*., 2021).

The plant PSR is governed by a complex regulatory network, with several components coordinating local (i.e., Pi sensing at the root tip) and long-distance signaling (Dissanayaka *et al*., 2021). During Pi-deprivation, proteins from the PHOSPHATE DEFICIENCY RESPONSE 2 (PDR2) and LOW PHOSPHATE ROOT (LPR) families interact to arrest primary root growth at the local level, whereas proteins containing a SYG1/Pho81/XPR1 (SPX) domain modulate the PSR by binding inositol pyrophosphates (Pi-signaling molecules responsive to Pi availability) (Ticconi *et al*., 2009; Jung *et al*., 2018). Notably, PHOSPHATE1 (PHO1), a xylem Pi-exporter responsible for root-to-shoot Pi translocation, as well as several intracellular PHTs, contain SPX domains and are thus sensitive to fluctuating Pi levels (Ham *et al*., 2018). Plants also contain PSR transcription factors (PHRs) which function as transcriptional activators of many PSI genes (Jung *et al*., 2018).

Despite identification of many PSI ‘marker’ genes, the complex signaling network that governs the PSR is extensive, with many components remaining undiscovered. Previous large-scale ‘Omic’ analyses focusing on plant Pi-starvation / resupply have focused on comparing genetic and / or proteomic changes at discrete time-points or within a specific tissue (reviewed in Zhou *et al*., 2022). The few studies that have investigated spatiotemporal PSI gene regulation reported that most are responsive following longer periods of Pi starvation / recovery, although there are some ‘early-response’ members including *TYPE 5 ACID PHOSPHATASE / PURPLE ACID PHOSPHATASE 17* (*ACP5/PAP17*) and *INDUCED BY PI STARVATION 1* (*IPS1*), which are reduced in transcript abundance by ∼30% within 1 h of Pi-resupply (Müller *et al*., 2004). Furthermore, Hammond *et al*., (2003) reported that the greatest proportion of both ‘early’- and ‘late’-responsive PSI genes belonged to metabolism, cell rescue (i.e. protection from oxidative stress and damage), and defense functional categories, with the proportion of genes involved in transcription, protein turnover, cellular communication and transport increasing in the ‘late-responsive’ category. Pi-sensing and gene expression also differ between tissues, with root transcriptional changes preceding changes in the shoots, with PSI genes being differentially expressed in different tissues, suggesting multiple regulatory systems governing the PSR (Wang *et al*., 2002; Müller *et al*., 2004).

Previously, we analyzed the impact of Pi resupply on the proteome of heterotrophic - Pi *Arabidopsis thaliana* (Arabidopsis) suspension cell cultures, identifying a combination of previously established and novel PSI proteins (Mehta *et al*., 2021). Here, we build on this work by distinguishing both temporal- and tissue-specific response elements of the PSR in Arabidopsis seedlings to gain a better understanding of its dynamic nature. With proteins representing the functional units of all eukaryotic cells, quantifying proteome-level changes is critical to understanding the molecular mechanisms underpinning plant responses. Using quantitative proteomic approaches, we define ‘early’ and ‘late’ shoot and root proteome responses to Pi-refeeding of -Pi Arabidopsis seedlings, revealing several combinations of distinct and overlapping spatiotemporal proteomic changes involved in diverse plant cell processes. Collectively, our global proteome analysis provides critical new insights into the plant PSR to aid in the translational development of crops that can better acclimate to -Pi conditions while assimilating applied Pi more efficiently.

## 2 MATERIALS AND METHODS

### 2.1 Plant Material and Treatment

*Arabidopsis thaliana* Col-0 (Arabidopsis) seedlings were grown in liquid culture by placing 5 mg of sterile, stratified seeds into 250 mL magenta boxes containing 50 mL of sterile 0.5x Murashige-Skoog medium (pH 5.7) containing 1% (w/v) sucrose and 0.2 mM Pi. Seedlings were cultivated under continuous illumination (100 μmol m^−2^ s^−1^) at 23 °C on an orbital platform (80 rpm). After 7 days, the medium was replaced with 50 mL of fresh medium containing 0 or 1.5 mM KH_2_PO_4_. Five days later the seedlings were either supplemented with 2 mM KH_2_PO_4_ or maintained under -Pi; 1 and 48 h later, the Pi resupplied and -Pi seedlings were harvested, rinsed with ultrapure water, and blotted dry. The shoots and roots arising from each liquid culture were rapidly separated before snap freezing in liquid N_2_ and lyophilizing. Dry weights of lyophilized shoots and roots were recorded.

### 2.2 Acid Phosphatase Extraction and Activity Assays, and Protein Concentration Determination

Lyophilized shoots or roots were ground to a powder using a mortar and pestle containing a small spatula of sand and homogenized (1:12; w/v for shoots, 1:30; w/v for roots) in 50 mM sodium acetate (pH 5.6) containing 1 mM EDTA, 1 mM dithiothreitol (DTT), 1 mM phenylmethylsulfonyl fluoride, 5 mM thiourea and 1% (w/v) insoluble poly(vinylpolypyrrolidone). Homogenates were centrifuged at 14,000 x *g* for 10 min at 4 °C and clarified extracts were stored on ice. APase activity was measured by coupling the hydrolysis of phosphoenolpyruvate (PEP) to pyruvate to the lactate dehydrogenase reaction and assaying at 23 °C by monitoring NADH oxidation at 340 nm using a Spectramax Plus Microplate spectrophotometer (Molecular Devices, Sunnyvale, CA, USA). The APase reaction mixture contained 50 mM sodium acetate (pH 5.6), 5 mM PEP, 10 mM MgCl_2_, 0.2 mM NADH, and 3 units ml^-1^ of desalted rabbit muscle lactate dehydrogenase in a final volume of 0.2 ml. All assays were initiated by the addition of clarified extract and corrected for any background NADH oxidation by omitting PEP from the reaction mixture. One unit (U) of activity is defined as the amount resulting in the use of 1 µmol min^-1^ of substrate. Protein concentrations were determined using the Bradford protein assay with bovine γ-globulin as the standard.

### 2.3 Tissue Processing, Proteome Extraction and Digestion

Lyophilized shoots and roots were ground using Geno/Grinder (SPEX SamplePrep) for 30 s at 1,200 rpm. Ground tissues were resuspended (1:2; w/v) in 50 mM Hepes-KOH (pH 8.0), 100 mM NaCl, and 2% (w/v) SDS. Protein extraction was performed by shaking samples at 1,000 rpm at 95 °C for 5 min using a tabletop shaker (Eppendorf ThermoMixer F2.0). The samples were then clarified by centrifugation at 20,000 x *g* for 10 min at 23 °C. The supernatant fluid was reduced with 10 mM DTT (Cat# D9779, Sigma) for 10 min at 23 °C, followed by alkylation with 30 mM iodoacetamide (Cat# I1149, Sigma) for 30 min at 23 °C in the dark. Peptides were then generated using a KingFisher APEX (Thermo Scientific) automated sample processing system as previously described without deviation (Mehta *et al*., 2022). Here, proteins were digested using sequencing grade trypsin (Cat# V5113; Promega) diluted in 50 mM triethylammonium bicarbonate buffer pH 8.5 (T7408; Sigma). Samples were acidified with trifluoroacetic acid (Cat# A117, Fisher) to a final concentration of 0.5% (v/v). Peptide desalting was performed as previously described (Mehta *et al*., 2022) using an OT-2 liquid handling robot (Opentrons Labworks Inc.) mounted with Omix C18 pipette tips (A5700310K; Agilent). Desalted peptides were dried by using a speedvac at 23 °C and stored at −80 °C prior to resuspension in 3.0% (v/v) acetonitrile / 0.1% (v/v) formic acid for mass spectrometer (MS) injection.

### 2.4 LC-MS/MS Analysis

Peptides were analyzed on a Fusion Lumos Orbitrap MS (Thermo Fisher Scientific). One microgram of peptide was injected per replicate using Easy-nLC 1200 system (Cat# LC140; Thermo Fisher Scientific) and an Acclaim PepMap 100 C18 trap column (Cat# 164750; Thermo Fisher Scientific) followed by analysis using a 50 cm Easy-Spray PepMap C18 analytical column (ES902; Thermo Fisher Scientific) warmed to 50 °C. Peptides were eluted into the MS at 300 nL min^-1^ using a segmented solvent B gradient of 0.1% (v/v) formic acid in 80% (v/v) acetonitrile (A998, Thermo Fisher Scientific) from 4 - 41% solvent B over 107 min as previously described (Mehta *et al*., 2022). A positive ion spray voltage of 2.3 kV was used with an ion transfer tube temperature of 300 °C and an RF lens setting of 40%.

### 2.5 BoxCar Data Independent Acquisition (DIA)

BoxCarDIA acquisition was performed as previously described, without deviation (Mehta *et al*., 2022). Here, full scan MS1 spectra (350–1400 *m*/*z*) were acquired with a resolution of 120,000 at 200 *m*/*z* with a normalized automatic gain control (AGC) target of 100% per BoxCar isolation window and IT set to automatic. MS1 spectra were acquired using two multiplexed targeted SIM scans of 10 BoxCar windows each. Fragment MS2 spectra were acquired at resolution of 30,000 across 28 38.5 *m*/*z* windows overlapping by 1 *m*/*z* using a dynamic maximum injection time and an AGC target value of 2000%. A minimum number of desired points across each peak set to 6. A summary methods file has been uploaded as part of the PRIDE data repository (see Data Availability).

### 2.6 Data Analysis

Downstream data analysis of all acquired proteomic data was performed using Spectronaut ver. 17 (Biognosys AG) with default settings as previously described (Mehta *et al*., 2022). All searches were made against a custom-made decoy (reversed) version of the Arabidopsis protein database from Araport 11 (27,533 protein encoding genes; ver. 2022-09-14). Briefly, search parameters included the following: trypsin digest permitting two missed cleavages, fixed modifications (carbamidomethyl (C)), variable modifications (oxidation (M)) and a peptide spectrum match, peptide and protein false discovery threshold of 0.01. Significantly changing differentially abundant proteins (DAPs) were determined and corrected for multiple comparisons (Bonferroni-corrected *p value* < 0.05; *q value;* Table S1).

### 2.7 Bioinformatics and Statistical Analysis

Gene Ontology (GO) enrichment analysis was done using the Database for Annotation, Visualization and Integrated Discovery (DAVID; https://davidbioinformatics.nih.gov/summary.jsp). All proteins quantified within the study were used as background and significance was determined based on biological processes where p-value <0.01. GO dot plots were made using R version 4.3.1 and R package *ggplot2*. Predicted subcellular localization of the significantly changing proteins was obtained with SUBA5 and the consensus subcellular localization predictor SUBAcon (https://suba.live/). Metabolic pathway analysis was performed using the Plant Metabolic Network (PMN; pathway tools version 28.0; https://pmn.plantcyc.org/). Association networks were generated using STRING-DB plugin (https://string-db.org/) in Cytoscape version 3.10.1 (https://cytoscape.org/) and enhancedGraphics version 1.5.5 (https://apps.cytoscape.org/apps/) was used to generate the CIRCOS surrounding the nodes. The minimum edge threshold was set 0.9 for the shoot network and 0.7 for root. Statistical analyses of all proteomic data used a Bonferroni-adjusted Student’s *t-*test to identify significantly changing protein groups (*q-value*) unless otherwise stated. All data were plotted in Graphpad Prism version 8 (https://www.graphpad.com/scientific-software/prism) and Microsoft Excel, and final figures were assembled using Affinity Designer software version 2.4.1 (https://affinity.serif.com/en-us/designer/).

## 3 RESULTS AND DISCUSSION

### 3.1 Biomass and APase activity determinations validate Pi status of Arabidopsis seedlings

Our Arabidopsis Pi starvation and re-feeding experiments were carried out using seedlings initially cultured for 7 days in medium containing 0.2 mM Pi. Seedlings were then transferred to Pi-deficient (20 µM) medium for 5 days, after which each seedling culture was directly supplemented with either no Pi (-Pi) or 2 mM Pi (Pi-resupplied).

Seedlings were then harvested after either 1 or 48 h, followed by shoot and root tissue lyophilization and processing for LC-MS/MS (Figure 1a). To validate the nutritional Pi status of the experimental samples, we measured biomass per seedling culture following lyophilization. At 48 h, the dry weight of Pi-resupplied seedling cultures was significantly (40%) greater than that of -Pi seedlings (Figure 1b). At 1 h, no significant differences were observed, although this is expected as changes in seedling biomass following Pi-resupply are typically observed after 24-48 h (Müller *et al*., 2004; Figure 1b).

**FIGURE 1.**
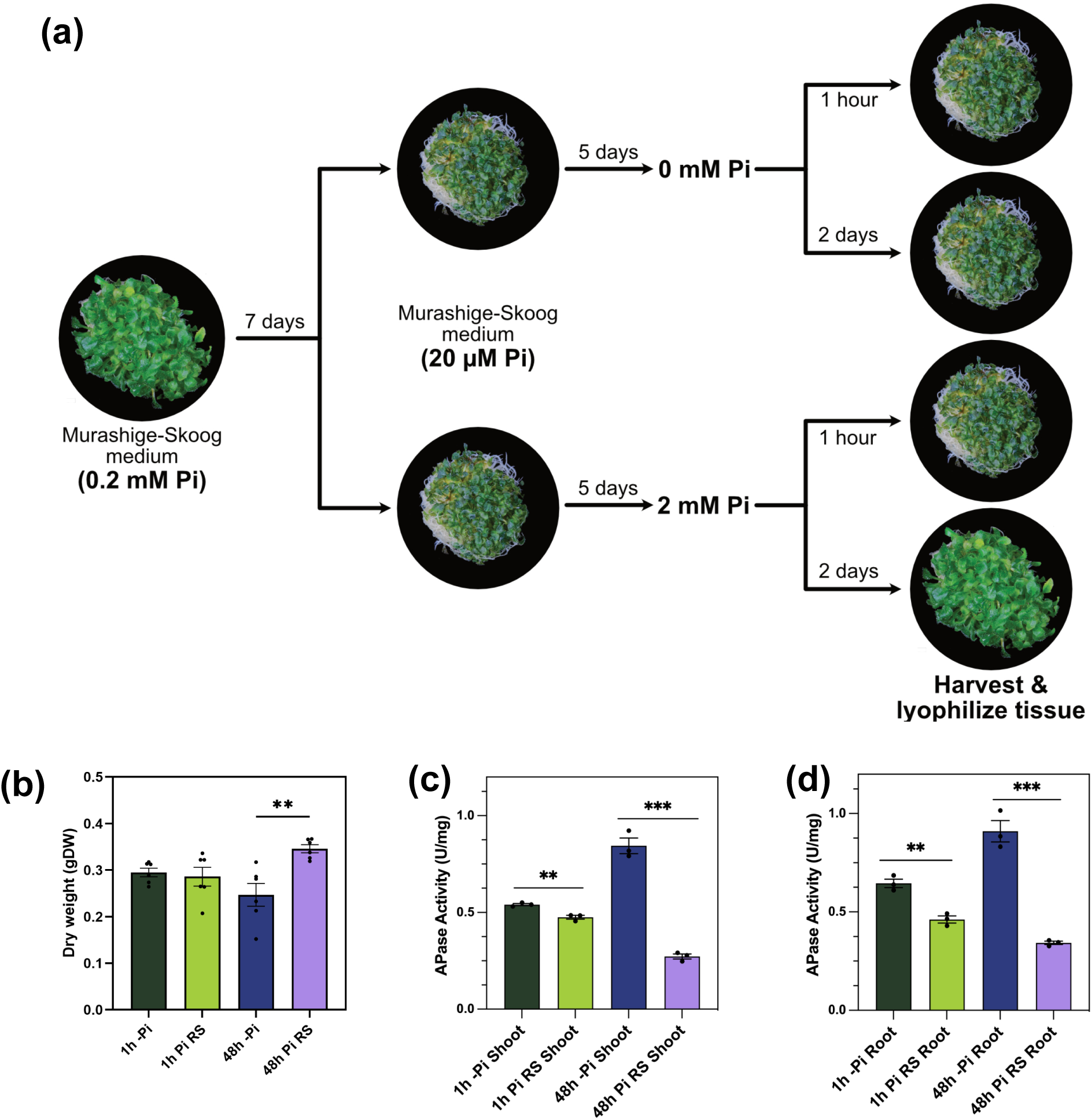
Sample preparation and validation. (a) Experimental workflow for preparation of 1 and 48 h Pi-starved (-Pi) and 2 mM Pi refed Arabidopsis seedlings. Images are examples of the -Pi and Pi-resupplied plants. (b) Seedling dry weight after 1 and 48h of Pi resupply represent means ±SE of n = 6 independent biological replicates. Significant differences were determined using the Student’s t-test (** P-value <0.05). Specific APase activities of the clarified protein extracts from (c) shoots and (d) roots after 1 and 48 h of Pi resupply represent means ±SE of *n* = 3 independent determinations using samples prepared from 3 biological replicates. Significant differences were determined using the Student’s t-test (***P-value < 0.05; ****P-value < 0.01).

A ubiquitous aspect of the plant PSR is the upregulation of secreted and vacuolar PAPs, which scavenge Pi from extracellular and expendable intracellular Po compounds (Veljanovski *et al*., 2006; Dissanayaka *et al*., 2021). We therefore assayed APase activities of clarified shoot and root extracts of -Pi plants, as well as following 1 and 48 h of Pi resupply. As expected, specific APase activity in the -Pi tissues was significantly greater than that of the Pi-resupplied tissues at both time-points (Figure 1c and 1d). We also compiled a list of over 20 PSR ‘marker proteins’ (i.e., those with documented evidence of their involvement in the Arabidopsis PSR) that were significantly reduced in abundance following Pi-resupply, to further validate the Pi nutritional status of our samples (Table 1). Among these were several PHTs, phospholipases, PAPs, and metabolic ‘bypass’ enzymes. In summary, our experimental model effectively induced Pi-starvation conditions, followed by Pi resupply-induced recovery.

**Table 1.**
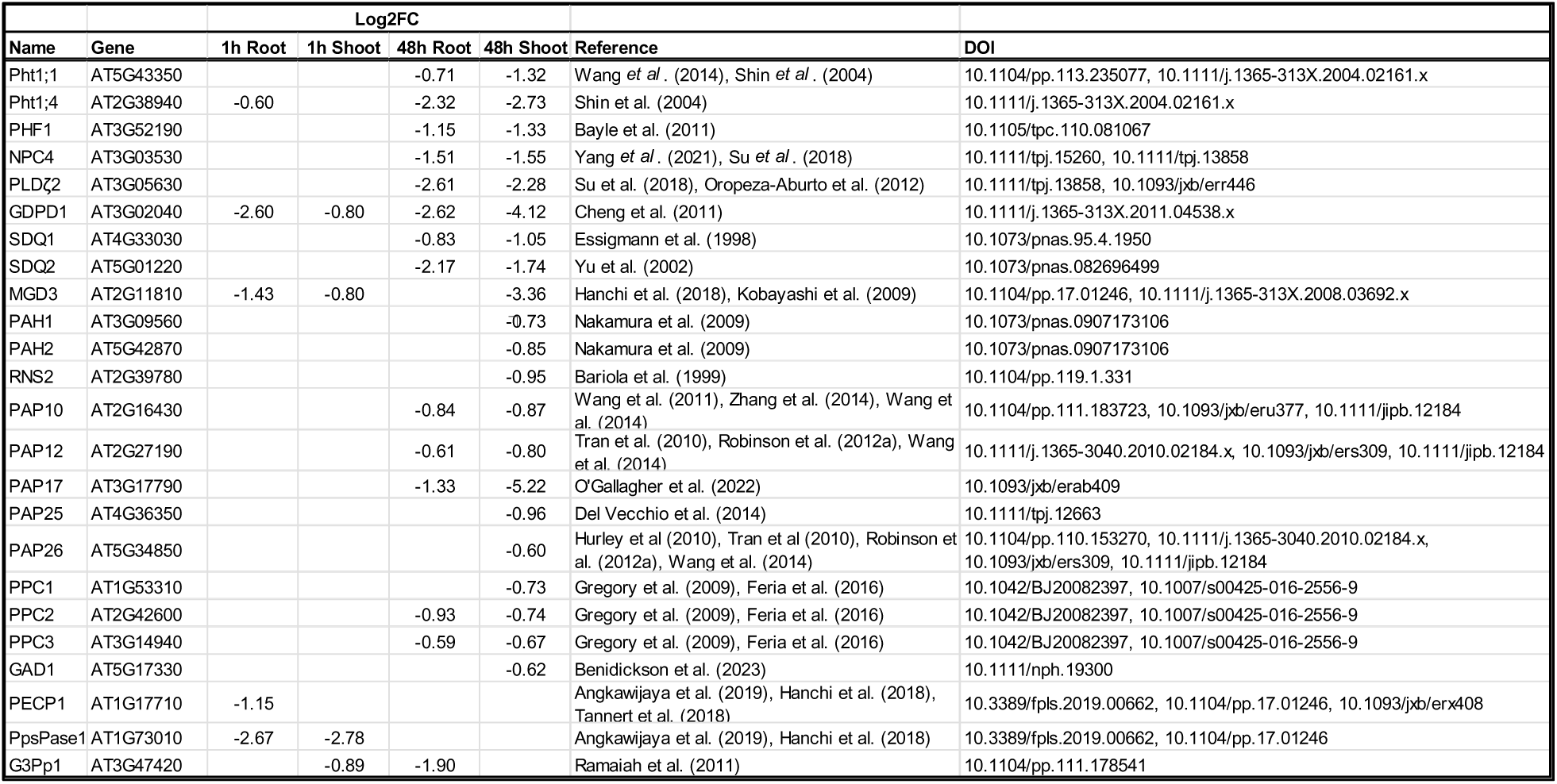
Established Pi starvation response proteins that were detected in our study.

### 3.2 Quantitative proteomic analysis reveals organ- and time-specific proteome changes upon Pi resupply

Quantitative LC-MS/MS was employed to analyze the shoot and root proteomes at 1 and 48 h following Pi-resupply. Across all treatments, ∼7,000 total proteins were quantified in the study, with ∼2,700 found to be significantly changing differentially abundant proteins (DAPs) (Log_2_FC > 0.58; *q* ≤ 0.05) (Table 2). Of these DAPs, 483 (root) and 608 (shoot) significantly changed in abundance 1 h following Pi-resupply, whereas 811 (root) and 1,527 (shoot) proteins significantly changed at the 48 h mark (*q*-value ≤ 0.05). In plotting the log-transformed fold changes of these significantly changing proteins, distinct trends emerge. At 1 h, the number of significantly changing DAPs increasing and decreasing in abundance is approximately equal in each direction (Figure 2a). However, at 48 h following Pi-resupply, 78% of root and shoot protein abundances change in tandem (i.e. either positively or negatively), whereas 22% change in opposing directions (Figure 2b), suggesting a temporal root-to-shoot sequence of proteome response. Furthermore, the overall magnitude of protein abundance change was much greater in the 48 h Pi-resupplied plants, highlighting the overall ‘late-responsive’ nature of recovery from Pi-starvation (Müller *et al*., 2004). At both time-points, there was a greater proportion of significantly changing proteins in the shoots relative to the roots, although this difference was magnified at the 48 h mark with nearly twice as many shoot proteins changing in abundance relative to roots (Figure 2c). Additionally, the predicted cellular localization of the significantly changing DAPs in shoots and roots at 1 and 48 h after Pi resupply resolved a greater percentage of plastid-localized proteins upregulated in the shoot at 1 h in comparison to 48 h shoot and both root time-points (Figure S1).

**FIGURE 2.**
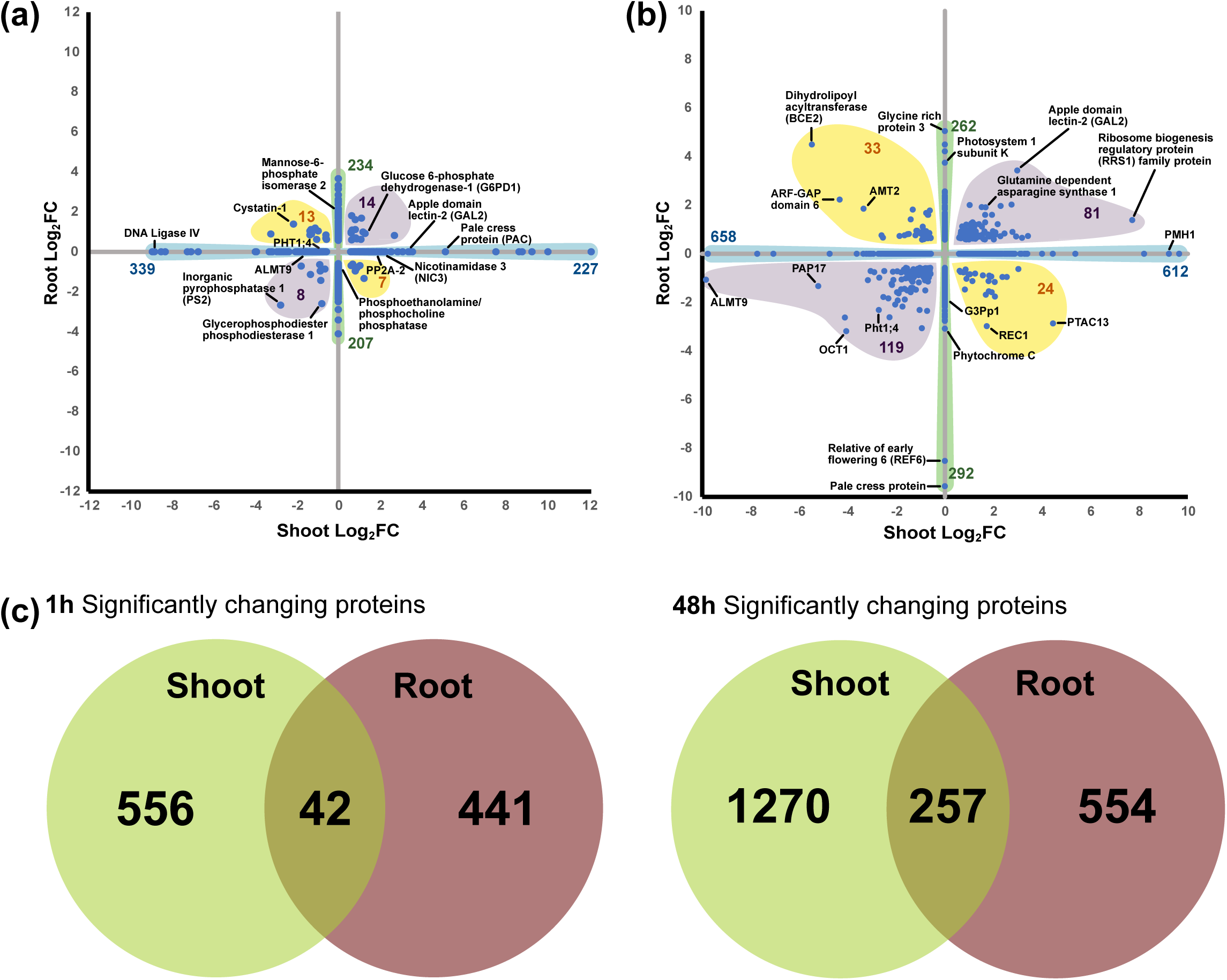
Distribution of significant shoot versus root proteome following Pi resupply of Arabidopsis seedlings. (a, b) Significantly changing shoot and root proteome (*q*-value < 0.05; *n* = 4; -0.58 > Log_2_ fold change (FC) > +0.58) in response to (a) 1 h or (b) 48 h of Pi resupply. Numbers denote the volume of significantly changed species on each axis or quadrant. Labeled proteins represent those with notable Log_2_FC in total abundance. (c) Venn diagrams depicting the overlap of significantly changing proteins common between or independent of 1 and 48 h of Pi resupply (*q*-value < 0.05; *n* = 4).

**Table 2.** Number of identified, quantified and significantly changing proteins.

|  | Identified | Quantified | Significantly Changing (1h) |  | Significantly Changing (48h) |  |
| --- | --- | --- | --- | --- | --- | --- |
|  |  |  | Root | Shoot | Root | Shoot |
| Proteins | 7068 | 5992 | 483 | 608 | 811 | 1527 |

### 3.3 The Pi responsive proteome: previously annotated and newly discovered Pi responsive proteins

#### 3.3.1 Pi sensing + transport

Plants acclimate to low Pi conditions through several molecular mechanisms, including the upregulation of PHTs that facilitate the uptake and transport of soil Pi to various plant tissues. Among these, plasma membrane-localized PHT1s are crucial for Pi uptake at the root-soil interface, with PHT1;1-1;4 contributing the most significantly of the nine PHT1 members found in Arabidopsis (Ayadi *et al*., 2015). Following Pi- resupply, we observed marked decreases in shoot and root PHT1;1 (AT5G43350) and PHT1;4 (AT2G38940) abundance (Figure S2), the latter of which was also corroborated in our previous study using Arabidopsis cell cultures (Mehta *et al*., 2021). In contrast, PHT5;3 (AT4G22990), one of the three Arabidopsis PHT5 members, was upregulated in roots following Pi-resupply (Figure S2). This upregulation suggests its role in sequestering excess intracellular Pi into vacuolar storage, as PHT5 proteins function as vacuolar Pi transporters that help to regulate cytoplasmic Pi levels (Liu *et al*., 2016). We also detected an increase in thylakoid membrane-localized PHT4;1 (AT2G29650) 1 h following Pi-resupply in the shoots (Figure S2). When grown under Pi-replete conditions, Arabidopsis *pht4;1* mutants demonstrated growth defects, likely due to PHT4;1’s role in chloroplast Pi compartmentation and ATP synthesis (Karlsson *et al*., 2015). These findings suggest PHT4;1 may also function as an ‘early-responsive’ PSR protein, facilitating rapid Pi delivery to the chloroplast stroma for ATP synthases and ultimately CO_2_ fixation following Pi-resupply.

Localization of Arabidopsis PHT1 proteins is partially coordinated by the PHOSPHATE TRANSPORTER TRAFFIC FACILITATOR (AT3G52190), which is induced under Pi starvation and mediates PHT1 exit from the endoplasmic reticulum (Bayle *et al*., 2011). Consistent with previous findings (Mehta *et al*., 2021), Pi transporter traffic facilitator-1 abundance declined in both shoots and roots 48 h following Pi-resupply (Figure S2). In contrast, we observed increased expression of PHO1;H3 (AT1G14040) (Figure S2), a PHO1 homologue involved in xylem loading and root-shoot Pi translocation, thus supporting shoot-targeted Pi allocation during recovery (Stefanovic *et al*., 2011). Further changes were also observed with GLYCEROL-3- PHOSPHATE PERMEASEs (G3Pp), which are additional ’Pi-responsive’ transporters that regulate G3P transport to maintain Pi homeostasis and redox balance (Table 1) (Ramaiah *et al*., 2011). Notably, *G3Pp1 (AT3G47420)* demonstrates an early and sustained pattern of upregulation following Pi-starvation (Misson *et al*., 2005; Ramaiah *et al*., 2011). Likewise, we found G3Pp1 was downregulated at early and late time- points following Pi-resupply (Figure S2), demonstrating its dynamic sensitivity to Pi-nutrition.

#### 3.3.2 Membrane lipid remodeling

A hallmark of the PSR is the upregulation of hydrolytic enzymes that liberate Pi from intra- and extracellular Pi-esters. Phospholipases are crucial to this process, as they facilitate Pi-scavenging and membrane lipid remodeling by facilitating the replacement of phospholipids with amphipathic galactolipids and sulfolipids (Dissanayaka *et al*., 2021). In our study, several phospholipases were significantly downregulated following Pi resupply in both shoots and roots, including NON-SPECIFIC PHOSPHOLIPASE C4 (NPC4; AT3G03530), PHOSPHOLIPASE 2 (PLC2; AT3G08510), PHOSPHOLIPASE Dζ2 (PLDζ2; AT3G05630), and glycerophosphodiester phosphodiesterase 1 (GDPD1; AT3G02040) (Figure S2) indicating a shift in lipid metabolism as the plants adjust to the restored Pi levels. NPC4 belongs to the NPC protein family, which hydrolyze phospholipids to produce diacylglycerol (DAG) and a phosphorylated head group (Yang *et al*., 2021). Upon Pi starvation, *npc4* knockout plants exhibit impaired phosphatidylcholine hydrolysis and reduced root hair elongation (Nakamura *et al*., 2005; Su *et al*., 2018; Yang *et al*., 2021). PLC2, although less well characterized in the context of Pi nutrition, has been implicated in stress signaling, including drought tolerance, pathogen defense, and auxin-regulated root growth (Georges *et al*., 2009; D’Ambrosio *et al*., 2017; Chen *et al*., 2019; van Hooren *et al*., 2024). Its downregulation suggests a broader role in Pi-associated stress signaling and lipid remode ling. PLDζ2 functions within the plant PSR to facilitate phosphatidylcholine hydrolysis to yield Pi (Su *et al*., 2018). Interestingly, the Arabidopsis *PLD*ζ*2* promoter contains an evolutionarily conserved Pi starvation-responsive transcriptional enhancer, which enhances root phospholipid degradation during Pi deficiency (Oropeza-Aburto *et al*., 2012). Additionally, *pld*ζ*2* knockout plants demonstrate reduced galactolipid replacement of membrane phospholipids during Pi-starvation (Su *et al*., 2018). GDPD1 hydrolyzes glycerophosphodiesters into glycerol-3-P and alcohol to liberate Pi and is transcriptionally induced during Pi starvation via PHR1-binding promoter elements in a time-dependent manner (Cheng *et al*., 2011). Our analysis determined that GDPD1 was strongly downregulated in both tissues and timepoints, complementing the results of Cheng *et al*. (2011) which supports its status as a dynamic PSR marker.

Sulfolipid replacement of phospholipids, an additional Pi-conserving strategy, requires the biosynthesis of sulfoquinovosyldiacylglycerol via two sulfolipid biosynthesis proteins, SULFOQUINOVOSYLDIACYLGLYCEROL 1 (SQD1; AT4G33030) and SDQ2 (AT5G01220). SQD1 converts UDP-glucose and sulfite into UDP-sulfoquinovose, a sulfolipid biosynthetic precursor, whereas SQD2 transfers this sulfoquinovose to DAG, forming sulfoquinovosyldiacylglycerol (Yu *et al*., 2002). We found both enzymes were downregulated 48 h following Pi resupply (Figure S2), consistent with previous findings that demonstrate their critical role in the PSR (Essigmann *et al*., 1998; Yu *et al*., 2002; Okazaki *et al*., 2009).

Plants also conserve Pi by substituting phospholipids with glycolipids such as monogalactosyldiacylglycerol (MGDG), synthesized by MGDG synthases (MGD1–3). MGD1 is the primary isozyme involved in bulk MGDG synthesis necessary for rapid membrane thylakoid expansion, whereas MGD2 and MGD3 play a more specialized role under Pi-limiting conditions, particularly in non-photosynthetic tissues (Awai *et al*., 2001; Kobayashi *et al*., 2009; Nitenberg *et al*., 2020). Our data revealed MGD1 (AT4G31780) was significantly downregulated in the shoots 48 h following Pi-resupply, whereas MGD3 (AT2G11810) was downregulated in both tissues at 1 h and in the shoots at 48 h (Figure S2). Although not previously linked to Pi-nutrition, MGD1 is sensitive to aluminum stress, likely due to its role(s) in lipid transport (Liu *et al*., 2020). As aluminum precipitates soil Pi and binds to negatively charged phospholipids in the plasma membrane, aluminum stress is often linked to Pi starvation; perhaps MGD1 has overlapping roles in mitigating aluminum toxicity and Pi starvation in plants (Panda and Matsumoto, 2009; Liu *et al*., 2020). MGD3 has been designated as a ‘fast responder’ to Pi-resupply, as demonstrated by its rapid transcriptional repression within 30 min of Pi- resupply in Arabidopsis roots (Hanchi *et al*., 2018). Furthermore, -Pi Arabidopsis *mgd3* mutants displayed drastic reductions in root glycolipid accumulation, fresh weight, root growth, and photosynthetic capabilities (Kobayashi *et al*., 2009). Our protein-level data reinforce MGD3’s dynamic and temporal involvement in plant Pi-recovery. Notably, neither MGD1 nor MGD3 were detected in Pi-resupplied Arabidopsis cell cultures (Mehta *et al*., 2021), highlighting the importance of a whole-plant tissue context.

Phosphatidate phosphatases (PAHs) hydrolyze membrane-bound phosphatidic acid for subsequent production of DAG (i.e., galactolipid precursor). Arabidopsis PAH1 (AT3G09560) and PAH2 (AT5G42870) are hypothesized to be a limiting step in Pi starvation-induced lipid remodeling; Arabidopsis *pah1pah2* double mutants demonstrated severe reductions in galactolipid production, elevated phospholipid content, impaired growth under N-deficiency, and heightened aluminum sensitivity under -Pi conditions (Nakamura *et al*., 2009; Kobayashi *et al*., 2013; Yoshitake *et al*., 2017). We detected a shoot-specific decrease in PAH1/2 abundance following 48 h of Pi-refeeding (Figure S2), suggesting a role in photosynthetic (i.e., chloroplast) membrane DAG production and subsequent MGD1 galactosylation for phospholipid membrane replacement (Nakamura *et al*., 2009). This is further evidenced by the lack of PAH1/2 identification in *Arabidopsis* cell cultures (Mehta *et al*., 2021), suggesting a Pi-starvation/resupply induced regulatory mechanism specific to autotrophic tissues.

Phosphoethanolamine / phosphocholine phosphatase 1 (PECP1; AT1G17710) and inorganic pyrophosphatase-1 (PPsPase1; AT1G73010) are well-established markers of Pi starvation that dephosphorylate phospholipid head groups to recycle Pi (Table 1) (Hanchi *et al*., 2018; Tannert *et al*., 2018; Angkawijaya *et al*., 2019). Following Pi- resupply, both phosphatases were strongly downregulated at 1 h (Figure S2), corroborating previous transcriptional evidence demonstrating a rapid (within 30 min) decrease in *PECP1* and *PPsPase1* mRNAs following Pi-replenishment (Hanchi *et al*., 2018). Given their dynamic expression in response to Pi availability, these phosphatases serve as sensitive ‘reporters’ of Pi starvation.

#### 3.3.3 Pi scavenging

Organic phosphorus (Po) compounds, such as nucleic acids derived from decaying biomatter, constitute 30–90% of the soil’s phosphorous pool (Dissanayaka *et al*., 2021). Likewise, endogenous nucleic acids account for nearly 50% of cellular Po of Pi-sufficient plants, with rRNA comprising 80% of this pool (Stigter and Plaxton, 2015). A hallmark of the plant PSR is the upregulation and root secretion of nucleases, ribonucleases (RNases), phosphodiesterases, and PAPs to liberate Pi from Po compounds (Dissanayaka *et al*., 2021). RNases are ubiquitous enzymes that maintain RNA degradation and turnover, and their Pi recycling abilities have been well-characterized in several plant species, including Arabidopsis, tomato, *Nicotiana alata*, rice, and snapdragon (Taylor *et al*., 1993; Köck *et al*., 1998; Bariola *et al*., 1999; Liang *et al*., 2002; Rojas *et al*., 2013; Gho *et al*., 2020). Arabidopsis ribonuclease 2 (RNS2) (AT2G39780) is constitutively expressed to mediate vacuolar RNA degradation and recycling, although its expression is further induced during Pi starvation or senescence to remobilize Pi (Taylor *et al*., 1993; Bariola *et al*., 1999; Liu *et al*., 2025). We found RNS2 was downregulated in shoots after 48 h of Pi-resupply (Figure S2), reinforcing its Pi-sensitivity. Interestingly, we found RNS3 (AT1G26820) was moderately downregulated in roots at 1 h and strongly downregulated in shoots at 48 h (Figure S2). RNS3, a secreted RNase, is typically induced by senescence rather than Pi starvation (Taylor *et al*., 1993; Bariola *et al*., 1994; Bariola *et al*., 1999). Given this discrepancy, perhaps RNS3 plays a more nuanced role with respect to Pi-recovery (rather than Pi-starvation) as the requirement for extracellular RNases rapidly decreases upon Pi- resupply. Furthermore, this discovery along with the lack of RNS2/3 detection in Arabidopsis cell cultures (Mehta *et al*., 2021) highlights the complex overlap between senescence and Pi-responsive signal transduction networks within various plant tissues (Dissanayaka *et al*., 2021).

APases hydrolyze Pi from organic Pi-monoesters and their upregulation is a distinctive feature of the plant PSR. PAPs are the largest class of plant APases, with the Arabidopsis genome encoding 29 different PAP isozymes that exhibit unique expression, localization, and metabolic patterns (Tran *et al*., 2010). Here, we found 13 PAPs were downregulated following Pi-resupply, and of these 13, PAP10, PAP12, PAP14, PAP17, PAP24, and PAP25 were also discovered in Arabidopsis cell cultures (Figure S2) (Tran *et al*., 2010; DelVecchio *et al*., 2014; Mehta *et al*., 2021; O’Gallagher *et al*., 2022). Previous literature has thoroughly characterized PAP10, PAP12, PAP17, PAP25, and PAP26 as the major Pi-responsive PAPs, as they are strongly upregulated at transcript and/or protein level during Pi deprivation (Del Pozo *et al*., 1999; Hurley *et al*., 2010; Tran *et al*., 2010; Wang *et al*., 2011; Robinson *et al*., 2012a; Robinson *et al*., 2012b; Wang *et al*., 2014; Del Vecchio *et al*., 2014; Zhang *et al*., 2014; Ghahremani *et al*., 2019; O’Gallagher *et al*., 2022). Furthermore, PAP12 and PAP26 are the main root-secreted and cell wall targeted APases that scavenge Pi from extracellular Pi-esters, while the dual-targeted AtPAP26 functions as the dominant vacuolar APase (Hurley *et al*., 2010; Robinson *et al*., 2012; Liu *et al*., 2025). Consistent with Mehta *et al*. (2021), PAP17 was the most downregulated PAP (in both tissues) following Pi resupply, underscoring its tight regulation during Pi-deprivation or leaf senescence (O’Gallagher *et al*., 2022). Interestingly, PAP27 was the only PAP downregulated at the 1 h resupply time-point, suggesting an ‘early-responsive’ role. Beyond Pi-starvation, PAP27 is also sensitive to pathogen infection and zinc deficiency (a condition with a well-established connection to Pi), hinting at a broader role in stress signaling (van de Mortel *et al*., 2006; Ascencio-Ibáñez *et al*., 2008; Khan *et al*., 2014).

Arabidopsis GALANTHUS NIVALIS AGGLUTININ-RELATED AND APPLE DOMAIN LECTIN-1 (GAL1), a dual-localized (cell wall and vacuole) mannose-binding lectin, interacts with a high mannose-glycoform of PAP26 to enhance its APase activity and stability to facilitate Pi scavenging during Pi starvation (Ghahremani *et al*., 2019; Ghahremani *et al*., 2019). Moreover, GAL1 was reported to be upregulated following Pi- but not phosphite-resupply in Arabidopsis cell cultures, suggesting a Pi-specific recovery response. While GAL1 is induced under Pi starvation (Ghahremani *et al*., 2019a), we found that its closest paralog, GAL2 (AT1G78860), was strongly upregulated following Pi resupply (Figure S2). This divergence in expression patterns could perhaps indicate these paralogs possess differential roles within the PSR, though more work is needed to explore GAL2’s unique role in Pi-recovery.

#### 3.3.4 Metabolic ‘bypass’ enzymes

Extended Pi starvation in plants depletes vacuolar Pi reserves, leading to reduced cytoplasmic Pi and Po levels (Dissanayaka *et al*., 2021). As several glycolytic enzymes rely on Pi-containing adenylates or Pi as a co-substrate, -Pi plants demonstrate significant metabolic and physiological impairments; however, they also possess remarkable metabolic redundancy, allowing them to bypass these steps through alternative adenylate and Pi-independent pathways. Plants upregulate several enzymes that utilize PPi as a co-substrate, which in contrast to Pi, ATP, and ADP is unaffected by Pi-deprivation (Duff *et al*., 1989). During Pi deprivation PEP carboxylase (PEPC) is simultaneously upregulated and activated by phosphorylation (Duff *et al.,* 1989, Gregory *et al*., 2009; Masakapalli *et al*., 2014; Mehta *et al*., 2019). In tandem with NAD-malate dehydrogenase and NAD-malic enzyme PEPC provides an ADP-independent bypass to pyruvate kinase, thus maintaining pyruvate availability for mitochondrial respiration while recycling Pi (a PEPC byproduct) under Pi-limiting conditions. Intermediate products of this bypass pathway include malate, which along with other carboxylates such as citrate is excreted by roots to mobilize Pi from the insoluble metal cation-Pi complexes in the soil (Dissanyanka *et al.,* 2021). Arabidopsis plant-type PEPC isozymes (PPC) 1–3 are Pi-responsive enzymes that are upregulated and activated by phosphorylation during Pi starvation (Gregory *et al*., 2009; Feria *et al*., 2016). Likewise, we found PPC2 (AT2G42600) and PPC3 (AT3G14940) were downregulated 48 h following Pi-resupply in both shoots and roots, while PPC1 (AT1G53310) decreased only in shoots (Figure S2). Interestingly, we found two isozymes of PEP carboxykinase (PCK1, AT4G37870; PCK2, AT5G65690), which catalyze the reverse reaction of PEPC, (i.e., oxaloacetate decarboxylation to PEP) were upregulated in shoots 48 h following Pi-resupply (Figure S2) (Rojas and Iglesias, 2023). To the best of our knowledge, no studies have characterized PCK as being Pi-responsive, although perhaps its Pi- resupply mediated upregulation suggests the point at which gluconeogenesis is reinitiated, converting stored carbon (e.g., malate) into PEP for sugar synthesis and restoration of anabolic growth post -Pi recovery (Dissanayaka *et al*., 2021).

Additionally, the oxidative pentose phosphate pathway (OPPP) is proposed to be a Pi recovery-inducible pathway due to its dual role in producing carbon precursors for macromolecule synthesis, and for restoring redox balance (i.e., NADPH production) when cell transitions from a stress-induced, quiescent metabolic state to a growth-promoting anabolic state (Mehta *et al*., 2021; Smith *et al*., 2024). Within this pathway, we found 6-phosphogluconolactonase-1 (AT1G13700) and glucose 6-P dehydrogenase 1 (G6PD1; AT5G35790) were upregulated following Pi-resupply (Figure S2). The latter finding corroborates that of Yin and Ashihara (2008), who reported rapid *G6PD1* upregulation 24 h following Pi refeeding of Arabidopsis cell cultures. Given that G6PD catalyzes the first committed step of the OPPP pathway, perhaps this ‘fast-responsive’ upregulation is required to jumpstart metabolic flux through this pathway to support the rapid resumption of cell growth that ensues upon Pi resupply.

We also found both glutamate decarboxylase (GAD) 1 (AT5G17330) and 4 (AT2G02010) were downregulated in the shoots 48 h following Pi-resupply (Figure S2).

GAD represents a branchpoint enzyme that mediates flux through the γ-aminobutyric acid (GABA) shunt, a tricarboxylic acid cycle bypass that helps drive mitochondrial respiration under stress conditions (Brown and Shelp, 2016). GAD1 has been hypothesized to upregulate GABA shunt flux during Pi-deprivation to circumvent thiamine diphosphate-dependent 2-oxoglutarate dehydrogenase activity, thus maintaining tricarboxylic acid cycle flux and cell respiration (Benidickson *et al*., 2023). This is evidenced by hyperphosphorylation of GAD1’s N-terminus following Pi-resupply of -Pi Arabidopsis cell cultures, phenotypic impairments in -Pi *atgad1* mutants compared to wild-type plants, as well as GAD1 transcript and polypeptide upregulation in -Pi shoots (Mehta *et al*., 2021; Benidickson *et al*., 2023). GABA is also known to modulate aluminum-activated malate transporter (ALMT) activity; notably, ALMT3 localizes to root hair membranes during Pi-starvation to mediate malate efflux for soil-Pi liberation (Ramesh *et al*., 2015; Gilliham and Tyerman, 2016; Maruyama *et al*., 2019). Interestingly, ALMT9 (AT3G18440), a malate-activated chloride channel involved in drought responses and stomatal aperture regulation, demonstrated the greatest change in abundance across our dataset, decreasing notably in both tissues following Pi resupply (Figure S2) (De Angeli *et al*., 2013). ALMT9 is also implicated in GABA-mediated stomatal signaling (Xu *et al*., 2021; Gilliham and Xu, 2022; Jaślan and De Angeli, 2022), suggesting a potential but unexplored role in Pi-related GABA signaling.

Pi deprived plants can also bypass the Pi- and ADP-dependent reactions of mitochondrial respiration (i.e., ATP synthases) via alternative oxidase (AOX). AOX minimizes the production of toxic reactive oxygen species by maintaining electron flow through the mitochondrial electron transport chain (from ubiquinone to O_2_) when electron flux through the energy-conserving cytochrome pathway is restricted by the low intracellular ADP and Pi levels that accompany prolonged Pi starvation (Parsons *et al*., 1999; Dissanayaka *et al*., 2021). In our study, AOX1A (AT3G22370) and AOX1D (AT1G32350) showed mixed responses to Pi-resupply (Figure S2). Previous work has demonstrated AOX1A is upregulated by Pi starvation and downregulated upon Pi resupply in Arabidopsis (Hammond *et al*., 2003; Vijayraghavan and Soole, 2010; Mehta *et al*., 2021). Interestingly, we observed divergent regulation of AOX1A between shoots and roots, along with a strong, early downregulation of AOX1D in the roots; these regulatory differences point to the complex, intricate metabolic flexibility that accompanies Pi starvation/resupply.

### 3.4 Gene Ontology and Network Analysis Identify Novel Biological Processes Impacted Upon Pi Resupply

In addition to known Pi-responsive markers, we were interested in identifying what other biological processes may be impacted at the proteome-level upon Pi resupply. To this end, we first performed a Gene Ontology (GO) enrichment analysis using the proteins that were significantly down- or upregulated upon Pi resupply at both 1 and 48 h time-points in root and shoot (Figure 3a and 3b). In correlation with the observed Pi markers mentioned above, the analysis identified enrichment of ‘Cellular Response to Phosphate Starvation’ (GO:0016036) among downregulated proteins in both shoots and roots at 48 h, as well as ‘Phospholipid Metabolic Process’ (GO:0006644) downregulated in root, and ‘Glycolipid Biosynthetic process’ (GO:0009247) downregulated in shoot, both at 48 h. We additionally noted enriched GO terms related to mRNA splicing among both root and shoot downregulated proteins, as well as a strong enrichment of GO terms relating to translation among upregulated proteins in shoot. To further define biological processes that may be impacted by the Pi resupply, we performed a STRING-DB association network analysis for both our shoot and root changing proteome (Figure 4). Protein association networks exhibiting significant changes in abundance under Pi- resupply treatments were created based on the StringDB database. As expected, both the shoot and root association networks showed a clear grouping of proteins primarily downregulated at 48 h with relation to Pi transport and metabolism as well as phospholipid metabolism. In addition, we observed nodes pertaining to the spliceosome, showing mostly downregulation at 48 h in root, with less of a clear pattern in shoot. Similarly, a grouping of proteins related to starch and sucrose metabolism showed a clear pattern of downregulation at 48 h in both shoot and root networks. One of the most striking patterns we observed among the association network was the clear upregulation of proteins involved in translation and ribosome biogenesis in shoots at 48 h of Pi resupply. Finally, we noted some differential regulation of photosynthesis-related proteins at the two time-points, with mostly upregulation at 1 h, followed primarily by downregulation at 48 h following Pi resupply. Next, we queried our data for metabolic pathways that may be significantly enriched, undertaking a plant metabolic pathway analysis (PMN; Table S2). In line with our GO and association network analyses, this strengthened the observation of changes in phospholipid and glycolipid metabolism, photosynthesis, carbohydrate metabolism and phenylpropanoid metabolism.

**FIGURE 3.**
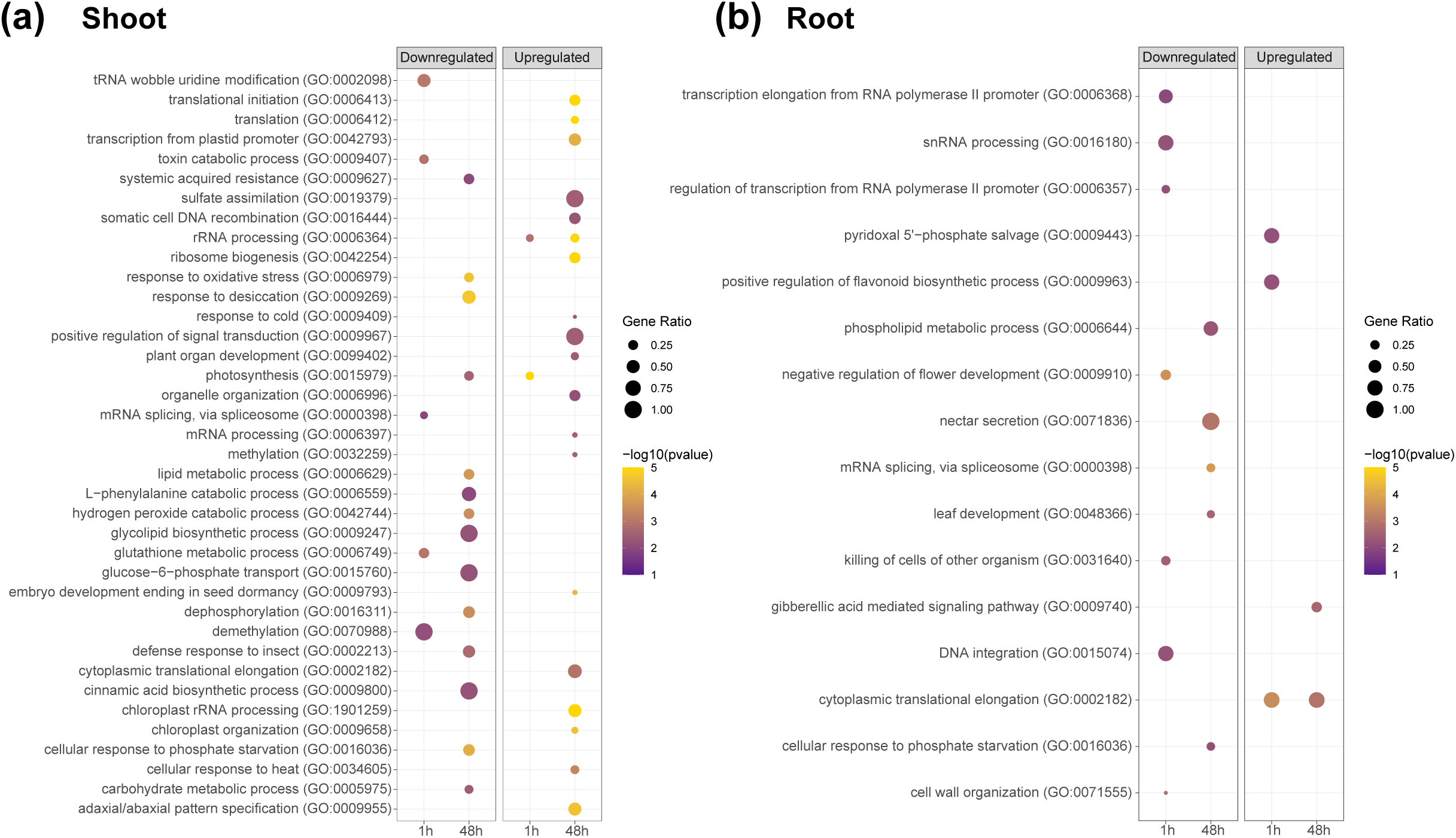
Gene Ontology (GO) enrichment analysis of shoot and root proteins increasing or decreasing upon Pi resupply. Dotplot representation of enriched biological process GO terms for significantly changing proteins (*q-*value <0.05; Log_2_FC>0.58 or <0.58) that are up- or down-regulated in (a) shoots or (b) roots following 1 or 48 h of Pi resupply. The dot size indicates the gene ratio (number of proteins seen under the condition/total quantified proteins in this study). The colour code of the dots represents log10 (*p*-value) on a gradient from low to high or purple to yellow.

**FIGURE 4.**
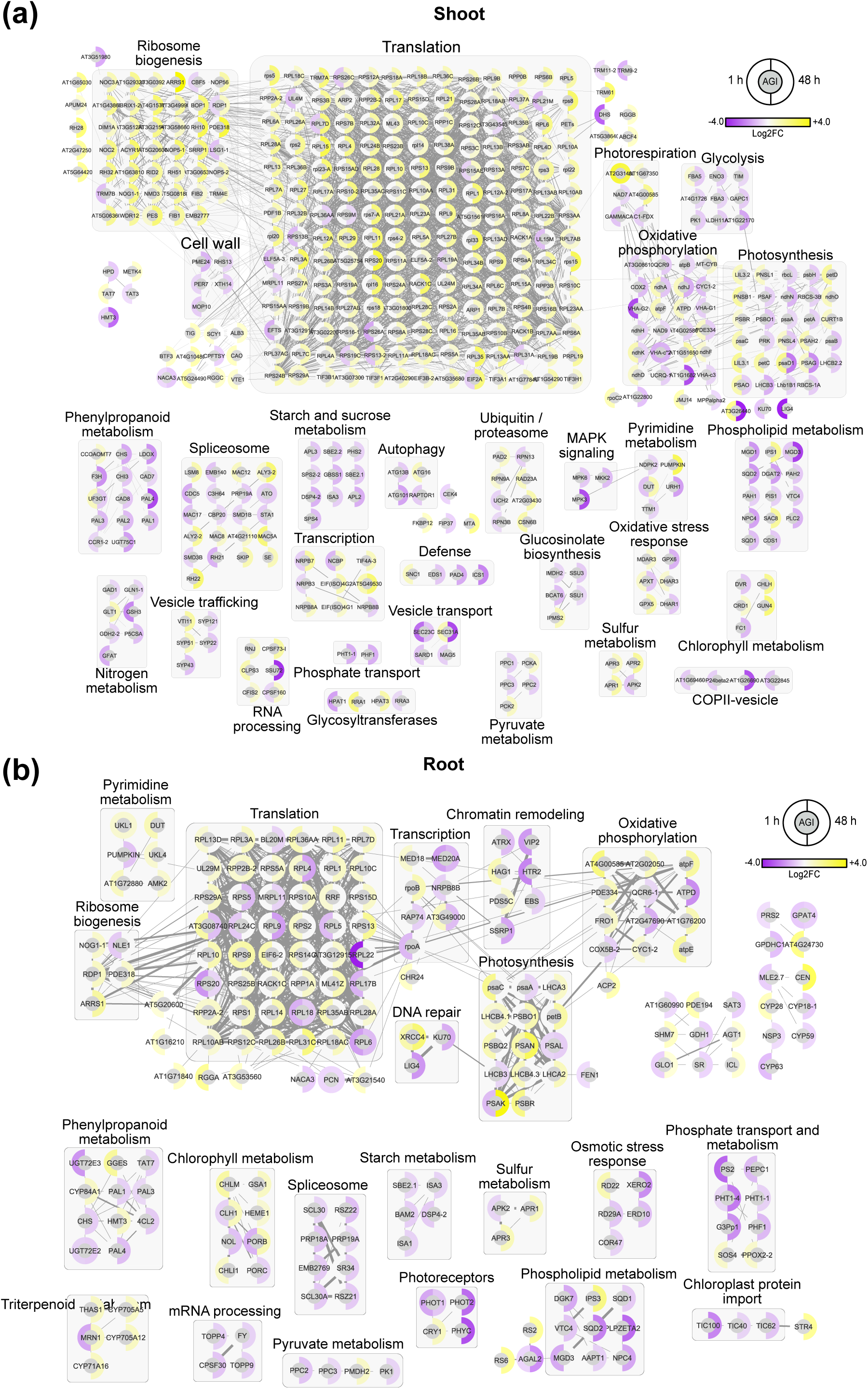
Association network analysis of proteins whose abundance changed following 1 and 48 h of Pi resupply. Association network analyses were performed using STRING-DB (ANOVA *p*-value <0.05) depicting significantly changing proteins following 1 and 48 h of Pi resupply in (a) shoots and (b) roots. Edge thickness indicates the strength of connection between nodes. Minimum edge threshold was set to 0.7 for roots and 0.9 for shoots. Node outer circles indicate the standardized relative Log_2_FC of the indicated protein in Pi resupply vs. -Pi. The scale of red to blue indicates relative increase and decrease in abundance, respectively. Grey circles encompass protein nodes involved in the same biological process.

#### 3.4.1 Translation and RNA splicing

Of all the categorized processes, the largest pool of significantly changing proteins (i.e., increased abundance) belonged to the ribosome biogenesis/translation network, particularly in the 48 h shoots (Figures 3 and 4). This large upregulation of protein synthesis machinery is expected, given Pi-starvation leads to an arrest in plant growth (and consequently down regulated protein synthesis), and a substantial proportion of cellular Pi is dedicated to rRNA (Wu *et al*., 2003; Veneklaas *et al*., 2012). In particular, the Arabidopsis ribosome biogenesis regulatory protein (RRS1) which directly modulates ribosome biogenesis was dramatically upregulated in the 48 h shoots (AT2G37990; 48 h shoot, +7.7169 Log_2_FC; 48 h root, +1.3912 Log_2_FC) (Choi *et al*., 2005).

Our data also identified mRNA splicing as downregulated in both tissues, though more so at 1 h in shoots and 48 h in roots (Figures 3 and 4). Several of these downregulated proteins were alternative splicing factors, particularly Ser/Arg (SR)-rich and SC35-like (SCL) proteins, which are involved in plant abiotic and nutritional stress responses (Chen *et al*., 2013; Zhang *et al*., 2014; Dong *et al*., 2018). In particular, several rice SR/SCL genes are involved in Pi homeostasis; two of which, SCL30/30a (Os12g38430/Os02g15310) and SR33 (Os07g0673500) have confirmed homology with Arabidopsis SCL30 (AT3G55460) and SR34 (AT1G02840), respectively, and were also downregulated in our study (AtSCL30; Shoot 1 h, -1.248 Log_2_FC, Shoot 48 h, -1.100 Log_2_FC, Root 48 h, -0.644 Log_2_FC; AtSR34; Root 48 h, -0.822 Log_2_FC) (Iida and Go, 2006; Dong *et al*., 2018; Gao *et al*., 2023). Furthermore, alternative splicing of REGULATOR OF LEAF INCLINATION 1 which modulates rice shoot architecture produces two protein isoforms that directly control PSR signaling pathways (Guo *et al*., 2022). Taken together, the PSI decrease in protein synthesis is likely a coordinated effect of decreased ribosome biogenesis coupled with increased alternative splicing.

#### 3.4.2 Carbohydrate Metabolism

Carbohydrate (i.e., starch and sucrose) metabolism enzymes in shoots was mostly downregulated at 48 h, although this response was varied in roots (Figures 3b and 4). In general, enzymes were upregulated that support anabolic growth, including G6PD1 involved in C-skeleton and nucleotide synthesis (discussed above) and ISOCITRATE LYASE (AT3G21720), a key enzyme of the glyoxylate cycle (Table S1). Several cell wall pectin/starch synthesis enzymes were also upregulated, including UDP-D-glucuronate 4-epimerase 1, 5, and 6 (Table S1), galacturonosyltransferase 1 (AT3G61130; root 48 h, +0.967 Log_2_FC), and PROTEIN TARGETING TO STARCH (AT5G39790; root 48 h +1.042 Log_2_FC). Contrary to this, enzymes related to C metabolism involved in the PSR were downregulated: sucrose-P synthase (SPS), a key enzyme involved in sucrose production, was downregulated in 48 h Pi-resupplied shoots and roots (SPS1F; AT5G20280; Root 48 h, -0.7756 Log_2_FC; SPS2F; AT5G11110; Shoot 48 h, -0.628 Log_2_FC, Root 48 h -0.8478 Log_2_FC; SPS4F; AT4G10120; Shoot 48 h, -1.096 Log_2_FC, Root 48 h, -0.8759). A characteristic early response of the PSR is the production of sucrose, which is a global regulator of Pi starvation signaling (Hammond and White, 2008; Lei *et al*., 2011). Furthermore, we found sucrose transporter 2 (SUC2) was downregulated in 48 h root tissues (AT2G02860; -0.8322 Log_2_FC). SUC2 facilitates root-sucrose transport, thus signaling the activation of Pi-responsive genes; disruption of the *SUC2* gene greatly inhibits plant PSR mechanisms (Lei *et al*., 2011).

#### 3.4.3 Phenylpropanoid metabolism

Under Pi starvation, phenylpropanoid biosynthesis is induced as they contribute to defense and structural modifications, including changes to root architecture (Luo *et al*., 2021; Wang *et al*., 2021). Correspondingly, we found phenylpropanoid biosynthesis was largely downregulated in shoot tissue following Pi resupply (Figure 4b). In particular, several enzymes involved in anthocyanin biosynthesis were downregulated, including leucoanthocyanidin dioxygenase (AT4G22880; Shoot 48 h -1.808 Log_2_FC), chalcone synthase (AT5G13930; Shoot 48 h -1.426 Log_2_FC), and flavonol synthase 5 (AT5G63600; Shoot 1 h -1.325 Log_2_FC). Leaf anthocyanin accumulation is a characteristic component of the plant PSR, as it protects against photoinhibitory damage during Pi-limited photosynthesis, whilst serving as metabolic markers of nutrient deficiency as seen by the classic purple pigmentation of -Pi plant leaves (Dissanayaka *et al*., 2021; Li *et al*., 2023). Previous evidence has identified leucoanthocyanidin dioxygenase as part of the plant PSR as its promoter contains PHR1 binding motifs (Liu *et al*., 2022). Chalcone synthase can also be linked to the PSR, as its expression is regulated by the Arabidopsis PRODUCTION OF ANTHOCYANIN PIGMENTS1 transcription factor; a Pi starvation-inducible regulator of anthocyanin biosynthesis (Li *et al*., 2016; Tao *et al*., 2024).

#### 3.4.4 Photosynthesis

In shoots, a major point of opposing regulation between the Pi-resupply timepoints was photosynthesis (i.e., photosynthetic proteins), which was generally upregulated at 1 h and downregulated at 48 h (Figures 3c and 4b; Supplemental Figure S1). As Pi is critical for ATP production and immediate restoration of metabolic homeostasis, we would expect a transient upregulation of photosynthetic machinery immediately following Pi supplementation (Carstensen *et al*., 2018). As sugar stores are replenished, there is a metabolic shift from photosynthetic recovery to other anabolic Pi-dependent processes, such as nucleotide synthesis, membrane rebuilding, and growth (Smith *et al*., 2024). Furthermore, prolonged leaf illumination without an adequate supply of Pi leads to photoinhibition and ROS production. Some specific photosynthetic proteins that followed this pattern were photosystem I subunits O, D1, and G (AT1G08380, AT4G02770, and AT1G55670 respectively), which were all upregulated at 1 h and downregulated at 48 h in shoots (Table S3). Interestingly, the dihydrolipoamide branched chain acyltransferase BCE2 had one of the most dramatically different regulatory patterns in shoots between timepoints (AT3G06850; 1 h, +3.35 Log_2_FC; 48 h, -5.48 Log_2_FC). BCE2 is a multifaceted protein that is induced during sucrose starvation, senescence/darkness, and in plants with impaired photosynthetic function (via mutation of the Thr78 phosphorylation site of rubisco activase) (Fujiki *et al*., 2005; Binder *et al*., 2007; Arias *et al*., 2014; Kim *et al*., 2019). Given sucrose accumulation and signaling are major regulators of the plant PSR, perhaps a rapid decrease in cytosolic sucrose levels following Pi-resupply (partially via direct allosteric inhibition of sucrose-P synthase by Pi) triggered an immediate upregulation of BCE2, followed by its downregulation at 48 h as sucrose and Pi levels stabilize (Doehlert and Huber, 1984; Lei *et al*., 2011). It would be of significance to explore BCE2’s putative role(s) in connecting Pi nutrition, sucrose signaling, and senescence.

In line with the regulatory pattern of photosynthetic proteins, two proteins involved in carbohydrate metabolism, fructose-bisphosphate aldolase 5 (FBA5; AT4G26530) and pullulanase-type limit dextrinase 1 (PU1; AT5G04360) were upregulated at 1 h then downregulated at 48 h (FBA5; shoot 1 h, +0.584 Log_2_FC, shoot 48 h, -0.795 Log_2_FC; PU1; root 1 h, +0.664 Log_2_FC, root 48 h, -2.34 Log_2_FC, shoot 48 h, -0.736 Log_2_FC). FBA5 is a cytosolic glycolytic enzyme that catalyzes fructose-1,6-P_2_/triose-P interconversion and is bound by calmodulin in a Ca^2+^-dependent manner (Symonds *et al*., 2024), whereas PU1 catalyzes the hydrolysis of the α-1,6 linkages of amylopectin during starch degradation, thus liberating glucose for glycolysis (Delatte *et al*., 2006). Upon replenished Pi stores, the immediate upregulation of these enzymes would simultaneously yield glucose and enhance glycolytic and respiratory flux for ATP production, followed by a shift towards anabolic growth at 48 h once energy reserves are restored.

In roots, the glycine-rich protein 3 (GRP3) was downregulated at 1 h, followed by a dramatic upregulation at 48 h (AT2G0552; Root 1 h, -1.22 Log_2_FC, Root 48 h, +5.05 Log_2_FC). GRP3 is a repressor of cell elongation, as demonstrated by increased root growth in *atgrp3* knockout plants, whose expression is induced by trehalose-6-P (T6P) (Schluepmann *et al*., 2004; Mangeon *et al*., 2016). Furthermore, GRP3 functions in the Arabidopsis wall-associated kinase 1-mediated aluminum signaling pathway, which may be connected to the PSR signaling pathway given the overlap between aluminum and Pi-stress response mechanisms (Sivaguru *et al*., 2003; Chen *et al*., 2022). The dynamic regulation of GRP3 likely mirrors cellular sucrose levels and T6P-mediated control of carbohydrate metabolism following Pi-resupply; elevated sucrose levels immediately following Pi-resupply result in increased conversion of G6P and UDP-Glucose to T6P by trehalose-6-P synthase, thus inhibiting sucrose nonfermenting1-related kinase1 (SnRK1) to promote growth (Müller *et al*., 2004; Schluepmann *et al*., 2004; Smeekens, 2015). After 48 h, the decline in sucrose levels facilitates T6P to trehalose conversion by trehalose-6-P phosphatase, thereby re-allocating resources from source to sink tissues likely through SnRK1’s phosphorylation and activation of C/S1 basic region-leucine zipper transcription factors (Smeekens, 2015).

## 4. Conclusions

Phosphate availability is a critical determinant of plant growth and development, influencing complex regulatory networks at molecular, cellular, and physiological levels. Our study resolved Pi-induced spatiotemporal changes to the Arabidopsis shoot and root proteomes, validating our current understanding and providing a number of novel insights into the dynamic nature of the PSR. We observed distinct and overlapping regulatory trends in shoots and roots at both early (1 h) and late (48 h) time-points. Classical PSR proteins (e.g., PHTs, PAPs, metabolic bypass enzymes, phospholipases) were generally downregulated following Pi resupply, although this downregulation varied between the two tissues and time-points highlighting a finely tuned system for Pi redistribution during recovery from Pi-starvation (Figure 5).

**FIGURE 5.**
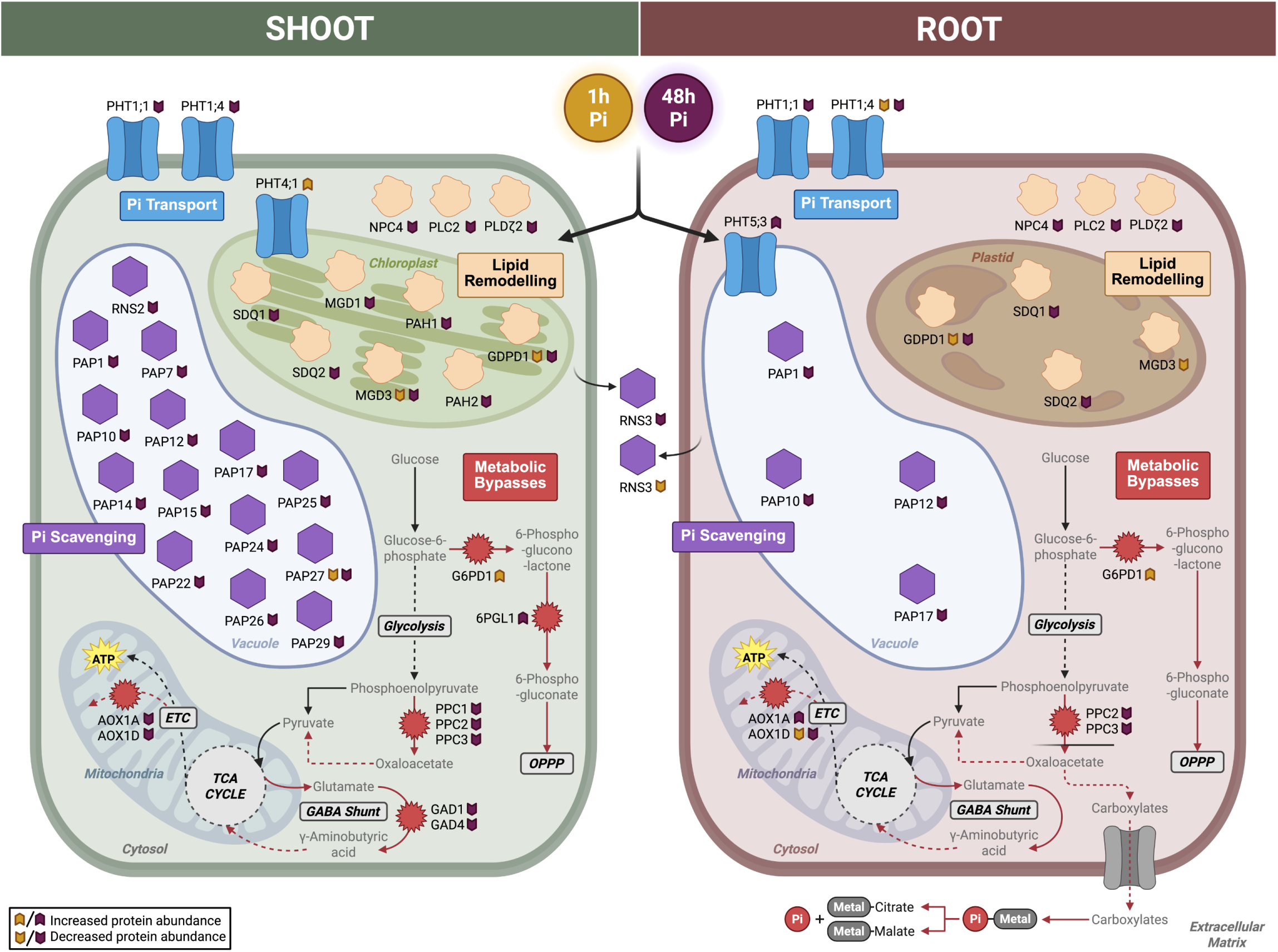
Summary of known proteomic changes observed following Pi refeeding in Arabidopsis shoots and roots. AOX, alternative oxidase; ETC, electron transport chain; GAD, glutamate decarboxylase; G6PD, glucose 6-P dehydrogenase; GDPD, glycerophosphodiester phosphodiesterase; MGD, monogalactosyldiacylglycerol synthase; NPC, non-specific phospholipase; OPPP, oxidative pentose phosphate pathway; PAH, phosphatidate phosphatase; PAP, purple acid phosphatase; 6PGL, 6- phosphogluconolactonase; PHT, Pi transporter; PL, phospholipase; PPC, plant type PEP carboxylase; RNS, ribonuclease; SDQ, sulfoquinovosyl diacylglycerol; TCA, tricarboxylic acid. Created in https://BioRender.com.

Overall, early responses predominantly involved metabolic adjustments to restore Pi pools via enhanced glycolysis and energy production, while late responses were characterized by metabolic shifts prioritizing anabolic processes including membrane remodeling and nucleotide production. This was especially notable in shoots, where the early upregulation of photosynthetic proteins at 1 h may reflect a transient need to restore photosynthetic capacity, followed by downregulation at 48 h that could help mitigate the risk of photoinhibition and ROS accumulation. Proteins involved in carbohydrate metabolism (e.g., FBA5, PU1, G6PD1) demonstrated a similar early upregulation followed by a delayed downregulation, reinforcing this trend. We also observed a dramatic upregulation of ribosomal proteins (i.e. translation machinery) at 48 h, and a concurrent downregulation of alternative splicing factors, some of which were implicated in plant abiotic and nutritional stress responses. Collectively, our study provides a comprehensive, proteome-centric framework of the plant PSR. These identifications, new and previously established, not only enhance our current understanding of plant nutritional stress responses but can also offer potential targets for biotechnological interventions aimed at improving crop Pi- acquisition and -use efficiency.

## ACKNOWLEDGEMENTS

Financial support was provided by research and equipment grants from the Natural Sciences and Engineering Research Council of Canada (NSERC) to WCP and RGU an NSERC CGS-D award to KHB, and a Canada Foundation for Innovation (CFI) award to RGU. The authors are grateful to Jack Moore of the Alberta Proteomics and Mass Spectrometry Facility for assistance with mass spectrometer operation and maintenance.

## AUTHOR CONTRIBUTIONS

MAS, LEG, KHB, DM - investigation; MAS, LEG, RGU - formal analysis; MAS, LEG, WCP, RGU - manuscript assembly and editing.

## DATA AVAILABILITY

Mass spectrometry data have been deposited to the ProteomeExchange Consortium (http://proteomecentral.proteomexchange.org) *via* the PRoteomics IDEntification Database (PRIDE; https://www.ebi.ac.uk/pride/) partner repository with the data set identifier PXD062427. **Username:****Password:** 4hoSHpIIofnU

## SUPPORTING INFORMATION

Table S1. All proteomics data for the manuscript. This includes: All quantified proteins, significantly changing proteins and Spectronaut v17 search parameter details.

Table S2. Enriched metabolic pathway analysis (PMN; https://www.plantcyc.org/databases/aracyc/8.0) using significantly changing proteins in shoots and roots following 1 and 48 h of Pi resupply.

Table S3. Differentially expressed significantly changing proteins

Figure S1. Subcellular localization of significantly changing proteins in Pi-resupplied *Arabidopsis* shoots and roots. *In silico* subcellular localization analysis of the significantly changing proteins that were down- or upregulated was performed using SUBAcon (https://suba.live/) for shoots and roots following 1 and 48 h of Pi resupply.

Figure S2. Summary of Pi responsive proteome. Heatmap of Pi-responsive proteome changes at 1 and 48 h after Pi resupply in shoots and roots relating to: (a) Pi sensing and transport, (b) membrane lipid remodeling, (c) Pi scavenging, and (d) and metabolic ‘bypass’ enzymes. Scale represents the Log_2_FC at the specified timepoint following Pi resupply where blue indicates downregulated and yellow indicates upregulated proteins.

## REFERENCES

1. Angkawijaya, A.E. et al. (2019) ‘Expression Profiles of 2 Phosphate Starvation-Inducible Phosphocholine/Phosphoethanolamine Phosphatases, PECP1 and PS2, in Arabidopsis’, Frontiers in Plant Science, 10. Available at: 10.3389/fpls.2019.00662.

2. Arias, M.C. et al. (2014) ‘From dusk till dawn: the Arabidopsis thaliana sugar starving responsive network’, Frontiers in Plant Science, 5. Available at: 10.3389/fpls.2014.00482.

3. Ascencio-Ibáñez, J.T. et al. (2008) ‘Global Analysis of Arabidopsis Gene Expression Uncovers a Complex Array of Changes Impacting Pathogen Response and Cell Cycle during Geminivirus Infection’, Plant Physiology, 148(1), pp. 436–454. Available at: 10.1104/pp.108.121038.

4. Awai, K. et al. (2001) ‘Two types of MGDG synthase genes, found widely in both 16:3 and 18:3 plants, differentially mediate galactolipid syntheses in photosynthetic and nonphotosynthetic tissues in Arabidopsis thaliana’, Proceedings of the National Academy of Sciences, 98(19), pp. 10960–10965. Available at: 10.1073/pnas.181331498.

5. Ayadi, A. et al. (2015) ‘Reducing the Genetic Redundancy of Arabidopsis PHOSPHATE TRANSPORTER1 Transporters to Study Phosphate Uptake and Signaling’, Plant Physiology, 167(4), pp. 1511–1526. Available at: 10.1104/pp.114.252338.

6. Bariola, P.A. et al. (1994) ‘The Arabidopsis ribonuclease gene RNS1 is tightly controlled in response to phosphate limitation’, The Plant Journal, 6(5), pp. 673–685. Available at: 10.1046/j.1365-313X.1994.6050673.x.

7. Bariola, P.A., MacIntosh, G.C. and Green, P.J. (1999) ‘Regulation of S-Like Ribonuclease Levels in Arabidopsis. Antisense Inhibition of RNS1 orRNS2 Elevates Anthocyanin Accumulation1’, Plant Physiology, 119(1), pp. 331–342. Available at: 10.1104/pp.119.1.331.

8. Bayle, V. et al. (2011) ‘Arabidopsis thaliana High-Affinity Phosphate Transporters Exhibit Multiple Levels of Posttranslational Regulation’, The Plant Cell, 23(4), pp. 1523–1535. Available at: 10.1105/tpc.110.081067.

9. Benidickson, K.H. et al. (2023) ‘Glutamate decarboxylase-1 is essential for efficient acclimation of Arabidopsis thaliana to nutritional phosphorus deprivation’, New Phytologist, 240(6), pp. 2372–2385. Available at: 10.1111/nph.19300.

10. Binder, S., Knill, T. and Schuster, J. (2007) ‘Branched-chain amino acid metabolism in higher plants’, Physiologia Plantarum, 129(1), pp. 68–78. Available at: 10.1111/j.1399-3054.2006.00800.x.

11. Blackwell, M., Darch, T. and Haslam, R. (2019) ‘Phosphorus use efficiency and fertilizers: future opportunities for improvements’, Frontiers of Agricultural Science and Engineering, 6(4), pp. 332–340. Available at: 10.15302/J-FASE-2019274.

13. Bown, A.W. and Shelp, B.J. (2016) ‘Plant GABA: Not Just a Metabolite’, Trends in Plant Science, 21(10), pp. 811–813. Available at: 10.1016/j.tplants.2016.08.001.

14. Carstensen, A. et al. (2018) ‘The Impacts of Phosphorus Deficiency on the Photosynthetic Electron Transport Chain’, Plant Physiology, 177(1), pp. 271–284. Available at: 10.1104/pp.17.01624.

15. Chen, T. et al. (2013) ‘A KH-Domain RNA-Binding Protein Interacts with FIERY2/CTD Phosphatase-Like 1 and Splicing Factors and Is Important for Pre-mRNA Splicing in Arabidopsis’, PLOS Genetics, 9(10), p. e1003875. Available at: 10.1371/journal.pgen.1003875.

16. Chen, W. et al. (2022) ‘Research Advances in the Mutual Mechanisms Regulating Response of Plant Roots to Phosphate Deficiency and Aluminum Toxicity’, International Journal of Molecular Sciences, 23(3), p. 1137. Available at: 10.3390/ijms23031137.

17. Chen, X. et al. (2019) ‘Phosphatidylinositol-specific phospholipase C2 functions in auxin- modulated root development’, *Plant*, Cell & Environment, 42(5), pp. 1441–1457. Available at: 10.1111/pce.13492.

18. Cheng, Y. et al. (2011) ‘Characterization of the Arabidopsis glycerophosphodiester phosphodiesterase (GDPD) family reveals a role of the plastid-localized AtGDPD1 in maintaining cellular phosphate homeostasis under phosphate starvation’, The Plant Journal, 66(5), pp. 781–795. Available at: 10.1111/j.1365-313X.2011.04538.x.

19. Choi, M.S. et al. (2005) ‘Isolation of a Calmodulin-binding Transcription Factor from Rice (Oryza sativa L.) *’, Journal of Biological Chemistry, 280(49), pp. 40820–40831. Available at: 10.1074/jbc.M504616200.

20. D’Ambrosio, J.M. et al. (2017) ‘Phospholipase C2 Affects MAMP-Triggered Immunity by Modulating ROS Production’, Plant Physiology, 175(2), pp. 970–981. Available at: 10.1104/pp.17.00173.

21. De Angeli, A. et al. (2013) ‘AtALMT9 is a malate-activated vacuolar chloride channel required for stomatal opening in Arabidopsis’, Nature Communications, 4(1), p. 1804. Available at: 10.1038/ncomms2815.

22. Del Pozo, J.C. et al. (1999) ‘A type 5 acid phosphatase gene from Arabidopsis thaliana is induced by phosphate starvation and by some other types of phosphate mobilising/oxidative stress conditions’, The Plant Journal, 19(5), pp. 579–589. Available at: 10.1046/j.1365-313X.1999.00562.x.

23. Del Vecchio, H.A. et al. (2014) ‘The cell wall-targeted purple acid phosphatase AtPAP25 is critical for acclimation of Arabidopsis thaliana to nutritional phosphorus deprivation’, The Plant Journal, 80(4), pp. 569–581. Available at: 10.1111/tpj.12663.

24. Delatte, T. et al. (2006) ‘Evidence for Distinct Mechanisms of Starch Granule Breakdown in Plants*’, Journal of Biological Chemistry, 281(17), pp. 12050–12059. Available at: 10.1074/jbc.M513661200.

25. Dissanayaka, D.M.S.B. et al. (2021) ‘Recent insights into the metabolic adaptations of phosphorus-deprived plants’, Journal of Experimental Botany, 72(2), pp. 199–223. Available at: 10.1093/jxb/eraa482.

26. Doehlert, D.C. and Huber, S.C. (1984) ‘Phosphate Inhibition of Spinach Leaf Sucrose Phosphate Synthase as Affected by Glucose-6-Phosphate and Phosphoglucoisomerase 1’, Plant Physiology, 76(1), pp. 250–253. Available at: 10.1104/pp.76.1.250.

27. Dong, C. et al. (2018) ‘Alternative Splicing Plays a Critical Role in Maintaining Mineral Nutrient Homeostasis in Rice (Oryza sativa)’, The Plant Cell, 30(10), pp. 2267–2285. Available at: 10.1105/tpc.18.00051.

28. Duff, S.M.G., Moorhead, G.B.G., Lefebvre, D.D., and Plaxton, W.C.P. (1989) ‘Phosphate starvation inducible ‘bypasses’ of adenylate and phosphate dependent glycolytic enzymes in *Brassica nigra* suspension cells’, Plant Physiology, 90, pp.1275–1278. 10.1104/pp.90.4.1275.

29. Essigmann, B. et al. (1998) ‘Phosphate availability affects the thylakoid lipid composition and the expression of SQD1, a6, #ne required for sulfolipid biosynthesis in Arabidopsis thaliana’, Proceedings of the National Academy of Sciences, 95(4), pp. 1950–1955. Available at: 10.1073/pnas.95.4.1950.

30. Feria, A.B. et al. (2016) ‘Phosphoenolpyruvate carboxylase (PEPC) and PEPC-kinase (PEPC-k) isoenzymes in Arabidopsis thaliana: role in control and abiotic stress conditions’, Planta, 244(4), pp. 901–913. Available at: 10.1007/s00425-016-2556-9.

31. Fujiki, Y. et al. (2005) ‘Response to Darkness of Late-Responsive Dark-Inducible Genes is Positively Regulated by Leaf Age and Negatively Regulated by Calmodulin-Antagonist-Sensitive Signalling in Arabidopsis thaliana’, Plant and Cell Physiology, 46(10), pp. 1741–1746. Available at: 10.1093/pcp/pci174.

32. Gao, R. et al. (2023) ‘Comprehensive study of serine/arginine-rich (SR) gene family in rice: characterization, evolution and expression analysis’, PeerJ, 11, p. e16193. Available at: 10.7717/peerj.16193.

33. Georges, F. et al. (2009) ‘Over-expression of Brassica napus phosphatidylinositol- phospholipase C2 in canola induces significant changes in gene expression and phytohormone distribution patterns, enhances drought tolerance and promotes early flowering and maturation’, *Plant*, Cell & Environment, 32(12), pp. 1664–1681. Available at: 10.1111/j.1365-3040.2009.02027.x.

34. Ghahremani, M., Park, J., et al. (2019a) ‘Lectin AtGAL1 interacts with high-mannose glycoform of the purple acid phosphatase AtPAP26 secreted by phosphate-starved Arabidopsis’, Plant, Cell & Environment, 42(4), pp. 1158–1166. Available at: 10.1111/pce.13463.

35. Ghahremani, M., Tran, H., et al. (2019b) ‘A glycoform of the secreted purple acid phosphatase AtPAP26 co-purifies with a mannose-binding lectin (AtGAL1) upregulated by phosphate-starved Arabidopsis’, Plant, Cell & Environment, 42(4), pp. 1139–1157. Available at: 10.1111/pce.13432.

36. Gho, Y.-S. et al. (2020) ‘Phosphate-Starvation-Inducible S-Like RNase Genes in Rice Are Involved in Phosphate Source Recycling by RNA Decay’, Frontiers in Plant Science, 11. Available at: 10.3389/fpls.2020.585561.

37. Gilliham, M. and Tyerman, S.D. (2016) ‘Linking Metabolism to Membrane Signaling: The GABA–Malate Connection’, Trends in Plant Science, 21(4), pp. 295–301. Available at: 10.1016/j.tplants.2015.11.011.

38. Gilliham, M. and Xu, B. (2022) ‘γ-Aminobutyric acid may directly or indirectly regulate Arabidopsis ALMT9’, Plant Physiology, 190(3), pp. 1570–1573. Available at: 10.1093/plphys/kiac399.

39. Gregory, A.L. et al. (2009) ‘In vivo regulatory phosphorylation of the phosphoenolpyruvate carboxylase AtPPC1 in phosphate-starved Arabidopsis thaliana’, Biochemical Journal, 420(1), pp. 57–65. Available at: 10.1042/BJ20082397.

40. Guo, M. et al. (2022) ‘Alternative splicing of REGULATOR OF LEAF INCLINATION 1 modulates phosphate starvation signaling and growth in plants’, The Plant Cell, 34(9), pp. 3319–3338. Available at: 10.1093/plcell/koac161.

41. Ham, B.-K. et al. (2018) ‘Insights into plant phosphate sensing and signaling’, Current Opinion in Biotechnology, 49, pp. 1–9. Available at: 10.1016/j.copbio.2017.07.005.

42. Hammond, J.P. et al. (2003) ‘Changes in Gene Expression in Arabidopsis Shoots during Phosphate Starvation and the Potential for Developing Smart Plants’, Plant Physiology, 132(2), pp. 578–596. Available at: 10.1104/pp.103.020941.

43. Hammond, J.P. and White, P.J. (2008) ‘Sucrose transport in the phloem: integrating root responses to phosphorus starvation’, Journal of Experimental Botany, 59(1), pp. 93–109. Available at: 10.1093/jxb/erm221.

44. Hanchi, M. et al. (2018) ‘The Phosphate Fast-Responsive Genes PECP1 and PPsPase1 Affect Phosphocholine and Phosphoethanolamine Content’, Plant Physiology, 176(4), pp. 2943–2962. Available at: 10.1104/pp.17.01246.

45. Hinsinger, P. (2001) ‘Bioavailability of soil inorganic P in the rhizosphere as affected by root- induced chemical changes: a review’, Plant and Soil, 237(2), pp. 173–195. Available at: 10.1023/A:1013351617532.

46. van Hooren, M. et al. (2024) ‘Ectopic Expression of Distinct PLC Genes Identifies “Compactness” as a Possible Architectural Shoot Strategy to Cope with Drought Stress’, Plant and Cell Physiology, 65(6), pp. 885–903. Available at: 10.1093/pcp/pcad123.

47. Hurley, B.A. et al. (2010) ‘The Dual-Targeted Purple Acid Phosphatase Isozyme AtPAP26 Is Essential for Efficient Acclimation of Arabidopsis to Nutritional Phosphate Deprivation’, Plant Physiology, 153(3), pp. 1112–1122. Available at: 10.1104/pp.110.153270.

48. Iida, K. and Go, M. (2006) ‘Survey of Conserved Alternative Splicing Events of mRNAs Encoding SR Proteins in Land Plants’, Molecular Biology and Evolution, 23(5), pp. 1085–1094. Available at: 10.1093/molbev/msj118.

49. Jaślan, J. and De Angeli, A. (2022) ‘Heterologous expression reveals that GABA does not directly inhibit the vacuolar anion channel AtALMT9’, Plant Physiology, 189(2), pp. 469–472. Available at: 10.1093/plphys/kiac132.

50. Jung, J.-Y. et al. (2018) ‘Control of plant phosphate homeostasis by inositol pyrophosphates and the SPX domain’, Current Opinion in Biotechnology, 49, pp. 156–162. Available at: 10.1016/j.copbio.2017.08.012.

51. Karlsson, P.M. et al. (2015) ‘The Arabidopsis thylakoid transporter PHT4;1 influences phosphate availability for ATP synthesis and plant growth’, The Plant Journal, 84(1), pp. 99–110. Available at: 10.1111/tpj.12962.

52. Khan, G.A. et al. (2014) ‘Coordination between zinc and phosphate homeostasis involves the transcription factor PHR1, the phosphate exporter PHO1, and its homologue PHO1;H3 in Arabidopsis’, Journal of Experimental Botany, 65(3), pp. 871–884. Available at: 10.1093/jxb/ert444.

53. Kim, S.Y. et al. (2019) ‘In vivo evidence for a regulatory role of phosphorylation of Arabidopsis Rubisco activase at the Thr78 site’, Proceedings of the National Academy of Sciences, 116(37), pp. 18723–18731. Available at: 10.1073/pnas.1812916116.

54. Kobayashi, K. et al. (2009) ‘Type-B monogalactosyldiacylglycerol synthases are involved in phosphate starvation-induced lipid remodeling, and are crucial for low-phosphate adaptation’, The Plant Journal, 57(2), pp. 322–331. Available at: 10.1111/j.1365-313X.2008.03692.x.

55. Kobayashi, Yasufumi et al. (2013) ‘Molecular and Physiological Analysis of Al3+ and H+ Rhizotoxicities at Moderately Acidic Conditions’, Plant Physiology, 163(1), pp. 180–192. Available at: 10.1104/pp.113.222893.

56. Köck, M. et al. (1998) ‘Extracellular administration of phosphate-sequestering metabolites induces ribonucleases in cultured tomato cells’, Planta, 204(3), pp. 404–407. Available at: 10.1007/s004250050273.

57. Lambers, H. and Plaxton, W.C. (2015) ‘Phosphorus: Back to the Roots’, in *Annual Plant Reviews Volume 48*. John Wiley & Sons, Ltd, pp. 1–22. Available at: 10.1002/9781118958841.ch1.

58. Lei, M. et al. (2011) ‘Genetic and Genomic Evidence That Sucrose Is a Global Regulator of Plant Responses to Phosphate Starvation in Arabidopsis’, Plant Physiology, 156(3), pp. 1116– 1130. Available at: 10.1104/pp.110.171736.

59. Leutert, M. et al. (2019) ‘R2-P2 rapid-robotic phosphoproteomics enables multidimensional cell signaling studies’, Molecular Systems Biology, 15(12), p. e9021. Available at: 10.15252/msb.20199021.

60. Li, H. et al. (2023) ‘Molecular mechanism of phosphorous signaling inducing anthocyanin accumulation in *Arabidopsis*’, Plant Physiology and Biochemistry, 196, pp. 121–129. Available at: 10.1016/j.plaphy.2023.01.029.

61. Li, S. et al. (2016) ‘MYB75 Phosphorylation by MPK4 Is Required for Light-Induced Anthocyanin Accumulation in Arabidopsis’, The Plant Cell, 28(11), pp. 2866–2883. Available at: 10.1105/tpc.16.00130.

62. Liang, L. et al. (2002) ‘*AhSL28*, a senescence- and phosphate starvation-induced S-like RNase gene in *Antirrhinum*’, Biochimica et Biophysica Acta (BBA) - Gene Structure and Expression, 1579(1), pp. 64–71. Available at: 10.1016/S0167-4781(02)00507-9.

63. Liu, C. et al. (2020) ‘Arabidopsis *mgd* mutants with reduced monogalactosyldiacylglycerol contents are hypersensitive to aluminium stress’, Ecotoxicology and Environmental Safety, 203, p. 110999. Available at: 10.1016/j.ecoenv.2020.110999.

64. Liu, T.-Y. et al. (2016) ‘Identification of plant vacuolar transporters mediating phosphate storage’, Nature Communications, 7(1), p. 11095. Available at: 10.1038/ncomms11095.

65. Liu, Z. et al. (2022) ‘PHR1 positively regulates phosphate starvation-induced anthocyanin accumulation through direct upregulation of genes F3’H and LDOX in Arabidopsis’, Planta, 256(2), p. 42. Available at: 10.1007/s00425-022-03952-w.

66. Liu, A.Y. et al. (2025_ ‘Vacuolar phosphatases are essential for efficient nucleotide salvage in Arabidopsis’, *Journal of Experimental Botany*, eraf168, 10.1093/jxb/eraf168

67. Luo, X. et al. (2021) ‘Phosphate deficiency enhances cotton resistance to *Verticillium dahliae* through activating jasmonic acid biosynthesis and phenylpropanoid pathway’, Plant Science, 302, p. 110724. Available at: 10.1016/j.plantsci.2020.110724.

68. Mangeon, A. et al. (2016) ‘AtGRP3 Is Implicated in Root Size and Aluminum Response Pathways in Arabidopsis’, PLOS ONE, 11(3), p. e0150583. Available at: 10.1371/journal.pone.0150583.

69. Maruyama, H. et al. (2019) ‘AtALMT3 is Involved in Malate Efflux Induced by Phosphorus Deficiency in Arabidopsis thaliana Root Hairs’, Plant and Cell Physiology, 60(1), pp. 107–115. Available at: 10.1093/pcp/pcy190.

70. Masakapalli, S.K. et al. (2014) ‘The metabolic flux phenotype of heterotrophic Arabidopsis cells reveals a flexible balance between the cytosolic and plastidic contributions to carbohydrate oxidation in response to phosphate limitation’, The Plant Journal, 78(6), pp. 964–977. Available at: 10.1111/tpj.12522.

71. Mehta, D. et al. (2021) ‘Phosphate and phosphite have a differential impact on the proteome and phosphoproteome of Arabidopsis suspension cell cultures’, The Plant Journal, 105(4), pp. 924–941. Available at: 10.1111/tpj.15078.

72. Mehta, D., Scandola, S. and Uhrig, R.G. (2022) ‘BoxCar and Library-Free Data-Independent Acquisition Substantially Improve the Depth, Range, and Completeness of Label-Free Quantitative Proteomics’, Analytical Chemistry, 94(2), pp. 793–802. Available at: 10.1021/acs.analchem.1c03338.

73. Misson, J. et al. (2005) ‘A genome-wide transcriptional analysis using Arabidopsis thaliana Affymetrix gene chips determined plant responses to phosphate deprivation’, Proceedings of the National Academy of Sciences, 102(33), pp. 11934–11939. Available at: 10.1073/pnas.0505266102.

74. van de Mortel, J.E. et al. (2006) ‘Large Expression Differences in Genes for Iron and Zinc Homeostasis, Stress Response, and Lignin Biosynthesis Distinguish Roots of Arabidopsis thaliana and the Related Metal Hyperaccumulator Thlaspi caerulescens’, Plant Physiology, 142(3), pp. 1127–1147. Available at: 10.1104/pp.106.082073.

75. Müller, R. et al. (2004) ‘Gene expression during recovery from phosphate starvation in roots and shoots of Arabidopsis thaliana’, Physiologia Plantarum, 122(2), pp. 233–243. Available at: 10.1111/j.1399-3054.2004.00394.x.

76. Nakamura, Y. et al. (2005) ‘A Novel Phosphatidylcholine-hydrolyzing Phospholipase C Induced by Phosphate Starvation in *Arabidopsis*\*’, Journal of Biological Chemistry, 280(9), pp. 7469– 7476. Available at: 10.1074/jbc.M408799200.

77. Nakamura, Y. et al. (2009) ‘Arabidopsis lipins mediate eukaryotic pathway of lipid metabolism and cope critically with phosphate starvation’, Proceedings of the National Academy of Sciences, 106(49), pp. 20978–20983. Available at: 10.1073/pnas.0907173106.

78. Nitenberg, M. et al. (2020) ‘Mechanism of activation of plant monogalactosyldiacylglycerol synthase 1 (MGD1) by phosphatidylglycerol’, Glycobiology, 30(6), pp. 396–406. Available at: 10.1093/glycob/cwz106.

79. O’Gallagher, B. et al. (2022) ‘Arabidopsis PAP17 is a dual-localized purple acid phosphatase up-regulated during phosphate deprivation, senescence, and oxidative stress’, Journal of Experimental Botany, 73(1), pp. 382–399. Available at: 10.1093/jxb/erab409.

80. Okazaki, Y. et al. (2009) ‘A Chloroplastic UDP-Glucose Pyrophosphorylase from Arabidopsis Is the Committed Enzyme for the First Step of Sulfolipid Biosynthesis’, The Plant Cell, 21(3), pp. 892–909. Available at: 10.1105/tpc.108.063925.

81. Oropeza-Aburto, A. et al. (2012) ‘Functional analysis of the Arabidopsis PLDZ2 promoter reveals an evolutionarily conserved low-Pi-responsive transcriptional enhancer element’, Journal of Experimental Botany, 63(5), pp. 2189–2202. Available at: 10.1093/jxb/err446.

82. Palma, D.A., Blumwald, E. and Plaxton, W.C. (2000) ‘Upregulation of vacuolar H+-translocating pyrophosphatase by phosphate starvation of Brassica napus (rapeseed) suspension cell cultures’, FEBS Letters, 486(2), pp. 155–158. Available at: 10.1016/S0014-5793(00)02266-3.

83. Panda, S.K., Baluška,František and and Matsumoto, H. (2009) ‘Aluminum stress signaling in plants’, Plant Signaling & Behavior, 4(7), pp. 592–597. Available at: 10.4161/psb.4.7.8903.

84. Parsons, H.L., Yip, J.Y.H. and Vanlerberghe, G.C. (1999) ‘Increased Respiratory Restriction during Phosphate-Limited Growth in Transgenic Tobacco Cells Lacking Alternative Oxidase1’, Plant Physiology, 121(4), pp. 1309–1320. Available at: 10.1104/pp.121.4.1309.

85. Ramaiah, M. et al. (2011) ‘Characterization of the Phosphate Starvation-Induced Glycerol-3- phosphate permease Gene Family in Arabidopsis’, Plant Physiology, 157(1), pp. 279–291. Available at: 10.1104/pp.111.178541.

86. Ramesh, S.A. et al. (2015) ‘GABA signalling modulates plant growth by directly regulating the activity of plant-specific anion transporters’, Nature Communications, 6(1), p. 7879. Available at: 10.1038/ncomms8879.

87. Robinson, W.D., Carson, I., et al. (2012a) ‘Eliminating the purple acid phosphatase AtPAP26 in Arabidopsis thaliana delays leaf senescence and impairs phosphorus remobilization’, New Phytologist, 196(4), pp. 1024–1029. Available at: 10.1111/nph.12006.

88. Robinson, W.D., Park, J., et al. (2012b) ‘The secreted purple acid phosphatase isozymes AtPAP12 and AtPAP26 play a pivotal role in extracellular phosphate-scavenging by Arabidopsis thaliana’, Journal of Experimental Botany, 63(18), pp. 6531–6542. Available at: 10.1093/jxb/ers309.

89. Rojas, B.E. and Iglesias, A.A. (2023) ‘Integrating multiple regulations on enzyme activity: the case of phosphoenolpyruvate carboxykinases’, AoB PLANTS, 15(4), p. plad053. Available at: 10.1093/aobpla/plad053.

90. Rojas, H.J., Roldán, J.A. and Goldraij, A. (2013) ‘NnSR1, a class III non-S-RNase constitutively expressed in styles, is induced in roots and stems under phosphate deficiency in Nicotiana alata’, Annals of Botany, 112(7), pp. 1351–1360. Available at: 10.1093/aob/mct207.

91. Schluepmann, H. et al. (2004) ‘Trehalose Mediated Growth Inhibition of Arabidopsis Seedlings Is Due to Trehalose-6-Phosphate Accumulation’, Plant Physiology, 135(2), pp. 879–890. Available at: 10.1104/pp.104.039503.

92. Shin, H., Shin, H.W., Dewbre, G.R., and Harrison, M.J. (2004). ‘Phosphate transport in *Arabidopsis:* Pht1;1 and Pht1;4 play a major role in phosphate acquisition from both low- and high-phosphate environments’, The Plant Journal, (39(4), pp. 629–642. Available at: 10.1111/j.1365-313X.2004.02161.x.

93. Sivaguru, M. et al. (2003) ‘Aluminum-Induced Gene Expression and Protein Localization of a Cell Wall-Associated Receptor Kinase in Arabidopsis’, Plant Physiology, 132(4), pp. 2256–2266. Available at: 10.1104/pp.103.022129.

94. Smeekens, S. (2015) ‘From Leaf to Kernel: Trehalose-6-Phosphate Signaling Moves Carbon in the Field’, Plant Physiology, 169(2), pp. 912–913. Available at: 10.1104/pp.15.01177.

95. Smith, M.A., Benidickson, K.H. and Plaxton, W.C. (2024) ‘In Vivo Phosphorylation of the Cytosolic Glucose-6-Phosphate Dehydrogenase Isozyme G6PD6 in Phosphate-Resupplied Arabidopsis thaliana Suspension Cells and Seedlings’, Plants, 13(1), p. 31. Available at: 10.3390/plants13010031.

96. Stefanovic, A. et al. (2011) ‘Over-expression of PHO1 in Arabidopsis leaves reveals its role in mediating phosphate efflux’, The Plant Journal, 66(4), pp. 689–699. Available at: 10.1111/j.1365-313X.2011.04532.x.

97. Stigter, K.A. and Plaxton, W.C. (2015) ‘Molecular Mechanisms of Phosphorus Metabolism and Transport during Leaf Senescence’, Plants, 4(4), pp. 773–798. Available at: 10.3390/plants4040773.

98. Su, Y. et al. (2018) ‘Different effects of phospholipase Dζ2 and non-specific phospholipase C4 on lipid remodeling and root hair growth in Arabidopsis response to phosphate deficiency’, The Plant Journal, 94(2), pp. 315–326. Available at: 10.1111/tpj.13858.

99. Symonds, K. et al. (2024) ‘Characterization of Arabidopsis aldolases AtFBA4, AtFBA5, and their inhibition by morin and interaction with calmodulin’, FEBS Letters, 598(15), pp. 1864–1876. Available at: 10.1002/1873-3468.14979.

100. Tannert, M. et al. (2018) ‘Pi starvation-dependent regulation of ethanolamine metabolism by phosphoethanolamine phosphatase PECP1 in Arabidopsis roots’, Journal of Experimental Botany, 69(3), pp. 467–481. Available at: 10.1093/jxb/erx408.

101. Tao, H. et al. (2024) ‘WRKY33 negatively regulates anthocyanin biosynthesis and cooperates with PHR1 to mediate acclimation to phosphate starvation’, Plant Communications, 5(5), p. 100821. Available at: 10.1016/j.xplc.2024.100821.

102. Taylor, C.B. et al. (1993) ‘RNS2: a senescence-associated RNase of Arabidopsis that diverged from the S-RNases before speciation.’, Proceedings of the National Academy of Sciences, 90(11), pp. 5118–5122. Available at: 10.1073/pnas.90.11.5118.

103. Theodorou, M.E. and Plaxton, W.C. (1994) ‘Induction of PPi-dependent phosphofructokinase by phosphate starvation in seedlings of Brassica nigra’, Plant, Cell & Environment, 17(3), pp. 287–294. Available at: 10.1111/j.1365-3040.1994.tb00294.x.

104. Ticconi, C.A. et al. (2009) ‘ER-resident proteins PDR2 and LPR1 mediate the developmental response of root meristems to phosphate availability’, Proceedings of the National Academy of Sciences, 106(33), pp. 14174–14179. Available at: 10.1073/pnas.0901778106.

105. Tran, H.T. et al. (2010) ‘Biochemical and molecular characterization of AtPAP12 and AtPAP26: the predominant purple acid phosphatase isozymes secreted by phosphate-starved Arabidopsis thaliana’, Plant, Cell & Environment, 33(11), pp. 1789–1803. Available at: 10.1111/j.1365-3040.2010.02184.x.

106. Tran, H.T., Hurley, B.A. and Plaxton, W.C. (2010) ‘Feeding hungry plants: The role of purple acid phosphatases in phosphate nutrition’, Plant Science, 179(1), pp. 14–27. Available at: 10.1016/j.plantsci.2010.04.005.

107. Veljanovski, V. et al. (2006) ‘Biochemical and Molecular Characterization of AtPAP26, a Vacuolar Purple Acid Phosphatase Up-Regulated in Phosphate-Deprived Arabidopsis Suspension Cells and Seedlings’, Plant Physiology, 142(3), pp. 1282–1293. Available at: 10.1104/pp.106.087171.

108. Veneklaas, E.J. et al. (2012) ‘Opportunities for improving phosphorus-use efficiency in crop plants’, New Phytologist, 195(2), pp. 306–320. Available at: 10.1111/j.1469-8137.2012.04190.x.

109. Vijayraghavan, V. and Soole, K. (2010) ‘Effect of short- and long-term phosphate stress on the non-phosphorylating pathway of mitochondrial electron transport in Arabidopsis thaliana’, Functional Plant Biology, 37(5), pp. 455–466. Available at: 10.1071/FP09206.

110. Wang, L. et al. (2011) ‘The Arabidopsis Purple Acid Phosphatase AtPAP10 Is Predominantly Associated with the Root Surface and Plays an Important Role in Plant Tolerance to Phosphate Limitation’, Plant Physiology, 157(3), pp. 1283–1299. Available at: 10.1104/pp.111.183723.

111. Wang, L. et al. (2014) ‘Comparative genetic analysis of Arabidopsis purple acid phosphatases AtPAP10, AtPAP12, and AtPAP26 provides new insights into their roles in plant adaptation to phosphate deprivation’, Journal of Integrative Plant Biology, 56(3), pp. 299–314. Available at: 10.1111/jipb.12184.

112. Wang, Yan et al. (2021) ‘Phosphate Uptake and Transport in Plants: An Elaborate Regulatory System’, Plant and Cell Physiology, 62(4), pp. 564–572. Available at: 10.1093/pcp/pcab011.

113. Wang, Yanting et al. (2021) ‘The role of strigolactones in P deficiency induced transcriptional changes in tomato roots’, BMC Plant Biology, 21(1), p. 349. Available at: 10.1186/s12870-021-03124-0.

114. Wang, Y.-H., Garvin, D.F. and Kochian, L.V. (2002) ‘Rapid Induction of Regulatory and Transporter Genes in Response to Phosphorus, Potassium, and Iron Deficiencies in Tomato Roots. Evidence for Cross Talk and Root/Rhizosphere-Mediated Signals’, Plant Physiology, 130(3), pp. 1361–1370. Available at: 10.1104/pp.008854.

115. Wu, P. et al. (2003) ‘Phosphate Starvation Triggers Distinct Alterations of Genome Expression in Arabidopsis Roots and Leaves’, Plant Physiology, 132(3), pp. 1260–1271. Available at: 10.1104/pp.103.021022.

116. Xu, B. et al. (2021) ‘GABA signalling modulates stomatal opening to enhance plant water use efficiency and drought resilience’, Nature Communications, 12(1), p. 1952. Available at: 10.1038/s41467-021-21694-3.

117. Yang, B. et al. (2021) ‘Acylation of non-specific phospholipase C4 determines its function in plant response to phosphate deficiency’, The Plant Journal, 106(6), pp. 1647–1659. Available at: 10.1111/tpj.15260.

118. Yin, Y. and Ashihar, H. (2008) ‘Expression of Glucose-6-phosphate Dehydrogenase and 6- Phosphogluconate Dehydrogenase Isoform Genes in Suspension-Cultured Arabidopsis thaliana Cells’, Zeitschrift für Naturforschung C, 63(9–10), pp. 713–720. Available at: 10.1515/znc-2008-9-1017.

119. Yoshitake, Y. et al. (2017) ‘Arabidopsis Phosphatidic Acid Phosphohydrolases Are Essential for Growth under Nitrogen-Depleted Conditions’, Frontiers in Plant Science, 8. Available at: 10.3389/fpls.2017.01847.

120. Yu, B., Xu, C. and Benning, C. (2002) ‘Arabidopsis disrupted in SQD2 encoding sulfolipid synthase is impaired in phosphate-limited growth’, Proceedings of the National Academy of Sciences, 99(8), pp. 5732–5737. Available at: 10.1073/pnas.082696499.

121. Zhang, W. et al. (2014) ‘Splicing factor SR34b mutation reduces cadmium tolerance in *Arabidopsis* by regulating iron-regulated transporter 1 gene’, Biochemical and Biophysical Research Communications, 455(3), pp. 312–317. Available at: 10.1016/j.bbrc.2014.11.017.

122. Zhang, Y. et al. (2014) ‘A major root-associated acid phosphatase in Arabidopsis, AtPAP10, is regulated by both local and systemic signals under phosphate starvation’, Journal of Experimental Botany, 65(22), pp. 6577–6588. Available at: 10.1093/jxb/eru377.

123. Zhou, M. et al. (2022) ‘Proteomic Analysis Dissects Molecular Mechanisms Underlying Plant Responses to Phosphorus Deficiency’, Cells, 11(4), p. 651. Available at: 10.3390/cells11040651.

