## Supplementary material for "Phosphate resupply differentially impacts the shoot and root proteomes of *Arabidopsis thaliana* seedlings": Supp Figure 1

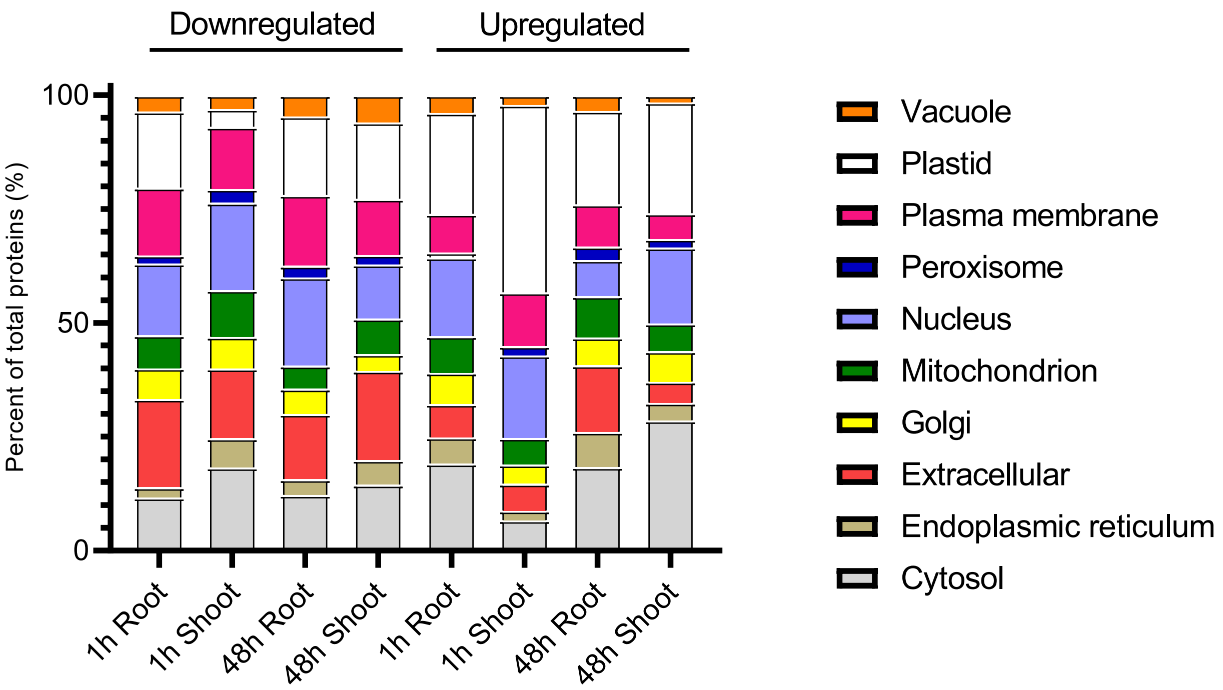


**Figure S1.** Subcellular localization of significantly changing proteins in Pi-resupplied *Arabidopsis* shoots and roots. *In silico* subcellular localization analysis of the significantly changing proteins that were down- or upregulated was performed using SUBAcon (https://suba.live/) for shoots and roots following 1 and 48 h of Pi resupply.
