## Supplementary material for "Phosphate resupply differentially impacts the shoot and root proteomes of *Arabidopsis thaliana* seedlings": Supp Figure 2

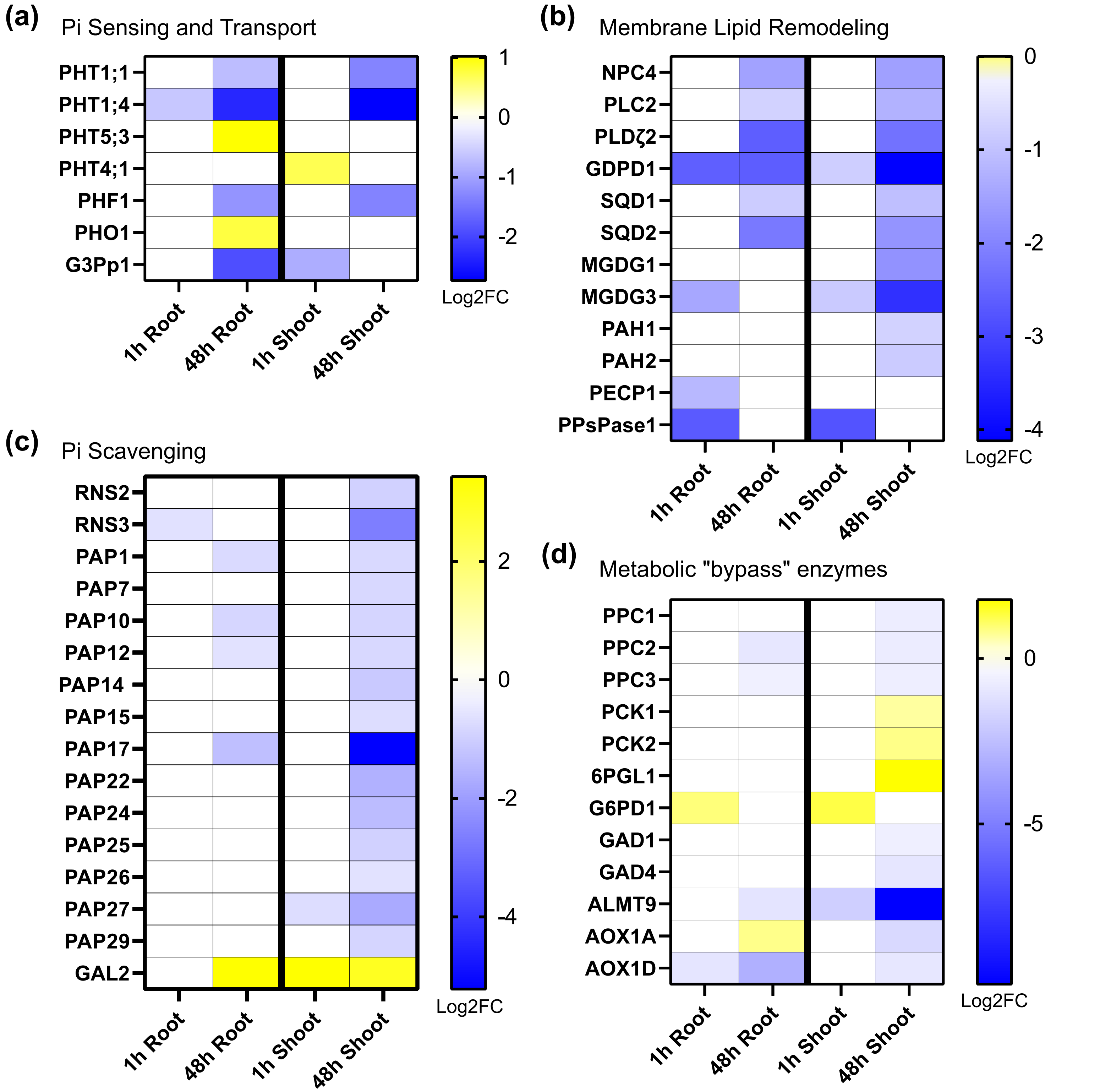


**Figure S2.** Summary of Pi responsive proteome. Heatmap of Pi-responsive proteome changes at 1 and 48 h after Pi resupply in shoots and roots relating to: (a) Pi sensing and transport, (b) membrane lipid remodeling, (c) Pi scavenging, and (d) and metabolic 'bypass' enzymes. Scale represents the Log2FC at the specified timepoint following Pi resupply where blue indicates downregulated and yellow indicates upregulated proteins.
